# Differential development of antibiotic resistance and virulence between *Acinetobacter* species

**DOI:** 10.1101/2023.12.01.569554

**Authors:** Elizabeth M. Darby, Robert A. Moran, Emma Holden, Theresa Morris, Freya Harrison, Barbara Clough, Ross S. McInnes, Ludwig Schneider, Eva M. Frickel, Mark A. Webber, Jessica M. A. Blair

## Abstract

The two species that account for most cases of *Acinetobacter*-associated bacteraemia in the UK are *Acinetobacter lwoffii*, often a commensal but also an emerging pathogen, and *A. baumannii*, a well-known antibiotic-resistant species. While these species both cause similar types of human infection and occupy the same niche, *A. lwoffii* (unlike *A. baumannii*) has thus far remained susceptible to antibiotics. Comparatively little is known about the biology of *A. lwoffii* and this is the largest study on it conducted to date, providing valuable insights into its behaviour and potential threat to human health.

This study aimed to explain the antibiotic susceptibility, virulence, and fundamental biological differences between these two species. The relative susceptibility of *A. lwoffii*, was explained as it encoded fewer antibiotic resistance and efflux pump genes than *A. baumannii* (9 and 30 respectively). While both species had markers of horizontal gene transfer, *A. lwoffii* encoded more DNA defence systems and harboured a far more restricted range of plasmids. Furthermore, *A. lwoffii* displayed a reduced ability to select for antibiotic resistance mutations, form biofilm and infect both *in vivo* and *in vitro* models of infection.

This study suggests that the emerging pathogen *A. lwoffii* has remained susceptible to antibiotics because mechanisms exist to make it highly selective about the DNA it acquires, and we hypothesise that the fact that it only harbours a single RND system restricts the ability to select for resistance mutations. This provides valuable insights into how development of resistance can be constrained in Gram negative bacteria.

**Importance:** *Acinetobacter lwoffii* is often a harmless commensal but is also an emerging pathogen and is the most common cause of *Acinetobacter*-derived blood stream infections in England and Wales. In contrast to the well-studied, and often highly drug resistant *A. baumannii*, *A. lwoffii* has remained susceptible to antibiotics. This study explains why this organism has not evolved resistance to antibiotics. These new insights are important to understand why and how some species develop antibiotic resistance, while others do not and could inform future novel treatment strategies.

## Introduction

*Acinetobacter* are Gram-negative, soil-dwelling, Gammaproteobacteria. Despite being typically found in the environment, some species within the genus also cause life-threatening human infections (1), the most clinically significant of these is *A. baumannii* which is often highly multidrug resistant (2, 3).

According to United Kingdom Health Security Agency (UKHSA), in England the most common cause of *Acinetobacter-*derived bacteraemia is *Acinetobacter lwoffii* followed by *A. baumannii* (30% and 21%, respectively) (4). *A. lwoffii* is found both in soil environments and as a common commensal of human skin (5). As well as causing bacteraemia in adults, *A. lwoffii* can cause a variety of infections, often in immunocompromised hosts and is a common cause of serious neonatal infections, which can lead to sepsis (6–9).

Both *A. lwoffii* and *A. baumannii* are found in hospitals and are resistant to desiccation, irradiation, and biocides (10, 11). However, *A. lwoffii* is generally antibiotic susceptible, in contrast to the multi-drug resistance displayed by *A. baumannii* (4). There are few studies aimed at understanding *A. lwoffii* and the reasons for its comparative sensitivity are not known.

We recently showed that the number of resistance nodulation division (RND) pumps present across the *Acinetobacter* genus varies and that *A. lwoffii* encodes fewer efflux pumps from the RND family than *A. baumannii* (1). These efflux pumps are important mediators of antibiotic resistance suggesting that their absence may contribute to the difference in susceptibility to antibiotics (12). RND pumps have also been implicated in virulence and biofilm formation (13, 14).

In this study we investigated the genomic and phenotypic differences between a range of *A. baumannii* and *A. lwoffii* strains (including clinical and type strains) to understand why two closely related species have such different responses to antibiotics. This study provides insight into the development of antibiotic resistance and differences in biology and virulence in two clinically important pathogens.

## Methods

### Strains used in this study

Reference strains of *A. baumannii* AYE and *A. lwoffii* NCTC 5867 were used. In addition, representative clinical and non-clinical strains were used in this study, listed in supplementary S1. All strains were cultured in lysogeny broth (LB) (Sigma) unless stated otherwise at 37°C.

### Measurement of the susceptibility of to antimicrobials

The minimum inhibitory concentration (MIC) of various antimicrobials to *A. baumannii* and *A. lwoffii* was determined using the agar dilution method (15) according to EUCAST (16). Antimicrobials tested included ampicillin (Sigma #A9393), cefotaxime (Fisher #10084487), chloramphenicol (Fisher #10368030), ciprofloxacin (Fisher #13531640), clindamycin (Generon #A10227), erythromycin (Fisher #10338080), fusidic acid (Sigma #F0881), gentamicin (Fisher #10224873), meropenem (TCI Chemicals #M2279), novobiocin (Fisher #15403619), rifampicin (Fisher #10533325) and tetracycline (Fisher #10460264).

### Biofilm formation and susceptibility

The ability of *A. baumannii* and *A. lwoffii* to establish monospecies biofilms and the susceptibility of these biofilms to different compounds was tested. The full methods can be found in supplementary S2.

### Whole genome sequence analysis

All available *A. lwoffii* and *A. baumannii* whole genome sequences were downloaded from NCBI (41 and 6,127 respectively) on 20/03/2022. In addition, laboratory strains of both *A. baumannii* (10) and *A. lwoffii* (8) were whole genome sequenced and assembled (MicrobesNG, UK). A list of strains sequenced in this study and their assembly accession numbers can be found in supplementary S3.

Quast (v.5.0.2) was used to quality check (QC) sequences and those with N50 values of <30,000 and >165 Ns per Kbp were removed (17). fastANI (v.1.31) was used to determine average nucleotide identity of *A. baumannii* sequences to *A. baumannii* AYE (CU459141.1) and *A. lwoffii* sequences to *A. lwoffii* 5867 (GCA_900444925.1) and only sequences >95.5% were kept (18). MASH (v.2.2.2) (19) was also performed to identify any duplicate assemblies which were then removed using a custom R script (https://github.com/C-Connor/MashDistDeReplication/blob/master/MashDistDeReplication.R). The final quality step was CheckM (v.1.1.3) (20), where sequences with >5% contamination and/or <95% completeness were removed. The final number of *A. baumannii* and *A. lwoffii* sequences was 4,809 and 38 respectively.

Assemblies were searched for antibiotic resistance genes (ARGs) (Comprehensive Antibiotic Resistance Database (21)), type IV pilus genes (‘twitching’ database using Ref. (22)), plasmid *rep* genes (database from Ref. (23)) and virulence and biofilm genes (‘vandb’ database using Ref. (24)) using ABRicate (v.0.8.13). The ‘twitching’ and ‘vandb’ databases can be found at: https://github.com/emd803/Gene-Databases/tree/main. Prophages were identified in a random 10 isolates of *A. baumannii* and *A. lwoffii* using PHASTER and DNA defence systems were searched for in all the genomes using DefenseFinder (v.1.0.9) (25, 26).

### Selection for resistance to meropenem, ciprofloxacin and gentamicin

To determine if *A. baumannii* (AB18) and *A. lwoffii* (AL28) could evolve resistance to three clinically relevant drugs, a selection experiment was set up, using strains clinically susceptible to all three selection antibiotics. Briefly, a single colony was inoculated into 5 mL of nutrient broth (Sigma) and a 1% transfer was passaged every 24 hours in increasing concentrations of each drug or without drug as a control. Populations from the terminal passage were spread onto LB agar and individual colonies were tested for their susceptibility to antibiotics listed above, as well as moxifloxacin (Sigma #PHR1542) and ethidium bromide (Fisher #10042120). Following selection, five colonies from parental strains AL28 and AB18 were subject to whole genome sequencing (MicrobesNG, UK) along with two colonies that had been passaged in nutrient broth only. Resulting sequences were compared to the appropriate parental strain.

Each whole genome sequence was confirmed to be from the species expected using ANI as above (>95%) and sequences were compared to both the ancestral strain and the cells passaged in nutrient broth only, using Snippy (v.4.6.0) to find sequence variants (27).

### Measurement of twitching motility and growth

A previously described crystal violet assay was used to measure twitching motility in *A. baumannii* and *A. lwoffii* (28). Additionally, growth in LB and human serum (Merck #H4522) was measured. Full methods in supplementary text S4.

### Scanning electron microscopy

Strains were grown overnight in LB, then diluted 1:50 for *A. baumannii* and 1:10 for *A. lwoffii* in LB because *A. lwoffii* grows to a lower final cell density than *A. baumannii*. Strains were grown to mid-log, washed with phosphate buffered saline (PBS) (Merck #D8537), and then resuspended in 2.5% glutaraldehyde (Sigma #354400) to fix. Cells were imaged on an Apreo 2 Scanning Electron Microscope (Thermo Fisher) at 5,000x, 10,000x and 25,000x magnification. Cell length analysis was performed in ImageJ (29) where the lengths of 100 randomly selected cells from each strain were measured.

### Virulence in the *Galleria mellonella* model

*Galleria mellonella* larvae were injected with 10^6^ bacterial cells as previously described (30) and the number of live/dead larvae were quantified across 7 days.

### Comparing the virulence in a macrophage cell line *in vitro*

Human monocyte THP-1 cell line (ATCC TIB-202) was cultured in Roswell Park Memorial Institute (RPMI) Medium with GlutaMAX (Thermo Fisher #61870-010) supplemented with 10% heat-inactivated fetal bovine serum (Life Technologies, #A5256701) at 37°C and 5% CO_2_. THP-1 monocytes were differentiated to macrophages with medium containing 50 ng/mL phorbol 12-myristate 13-acetate (PMA) (Sigma #P1585) for 3 days. Cells were then left to rest for 2 days by replacing the differentiation medium with complete medium without PMA. Macrophages were infected as previously described (31), with a multiplicity of infection (MOI) of 100. Extracellular bacteria were killed after 2 hours using gentamicin at either 100 μg/mL or at 1 mg/mL for AB05. Association, invasion, and proliferation (after 6 hours) were quantified. Association was determined by subtracting the number of intracellular bacteria (invasion) from the total number of bacteria associated with macrophages (and within macrophages).

## Results

### *A. lwoffii* is more susceptible to a broad range of antibiotics than *A. baumannii*

Data from the UKHSA shows that *A. lwoffii* isolated from patients in England were more susceptible than *A. baumannii* to gentamicin, ciprofloxacin, meropenem and colistin (4). Therefore, we sought to determine if the same was true in our diverse strain collection of strains for a range of antibiotics from different drug classes (Table 1). MICs were higher for *A. baumannii* than for *A. lwoffii* for all compounds tested. EUCAST resistance breakpoints were only available for ciprofloxacin (>1 μg/mL), meropenem (>2 μg/mL) and gentamicin (>4 μg/mL) (16). *A. lwoffii* was clinically susceptible in all instances, whereas for *A. baumannii*, all but one isolate was resistant to ciprofloxacin, three of six strains were resistant to meropenem, and all were resistant to gentamicin.

**Table 1.** MIC values for *A. baumannii* and *A. lwoffii* (μg/mL)

| Strain | AMP | CEF | CHL | CIP | CLI | ERY | FUS | GEN | MER | NOV | RIF | TET |
| --- | --- | --- | --- | --- | --- | --- | --- | --- | --- | --- | --- | --- |
| AB05 | 512 | >32 | 256 | >32 | 128 | 32 | 128 | 1024 | 0.5 | 16 | 16 | 128 |
| **AB18** | 64 | 16 | 128 | 1 | 64 | 16 | 128 | 4 | 0.25 | 8 | 4 | 1 |
| **AB19** | 8 | 32 | 128 | >32 | 64 | 32 | 128 | 16 | 0.25 | 8 | 4 | 8 |
| **AB20** | 32 | 16 | 128 | 2 | 64 | 64 | 64 | 4 | >16 | 16 | 4 | >128 |
| **AB25** | 1024 | >32 | 256 | >32 | 128 | 64 | 128 | 128 | 8 | 32 | 4 | >128 |
| **AB27** | 1024 | 16 | 128 | >32 | 32 | 8 | 64 | 1024 | 16 | 8 | 4 | >128 |
| **AL04** | <1 | 2 | 2 | 0.06 | 2 | <0.5 | 16 | <2 | 0.03 | 8 | 0.5 | 0.5 |
| **AL28** | <1 | 1 | 1 | 0.06 | 1 | 0.5 | 4 | <2 | 0.12 | 4 | 0.5 | 0.25 |
| **AL29** | <1 | 2 | 1 | 0.06 | 2 | 0.5 | 8 | <2 | 0.03 | 8 | 0.5 | 0.25 |
| **AL32** | <1 | 1 | 1 | 0.03 | 4 | 0.5 | 8 | <2 | 0.12 | 8 | 0.5 | 0.25 |
| **AL33** | <1 | 1 | 1 | 0.06 | 4 | 0.5 | 8 | <2 | 0.12 | 8 | 0.5 | 0.25 |

AMP - ampicillin, CEF - cefotaxime, CHL - chloramphenicol, CIP - ciprofloxacin, CLI - clindamycin, ERY - erythromycin, FUS - fusidic acid, GEN - gentamicin, MER - meropenem, NOV - novobiocin, RIF - rifampicin, TET - tetracycline

### *A. lwoffii* carries fewer antibiotic resistance genes (ARGs) than *A. baumannii*

To explain the differences in antibiotic sensitivity between *A. lwoffii* and *A. baumannii,* whole genome sequences were searched for the presence of ARGs using the CARD database. Following QC there were 4,809 *A. baumannii* and 38 *A. lwoffii* genome sequences. Across the *A. lwoffii* genomes 40 different ARGs were found whilst 333 different ARGs were detected across *A. baumannii.* Due to the lack of available sequences for *A. lwoffii,* to quantitatively compare the presence of ARGs between the two species, a random permutation was conducted, which subsampled 38 sequences (the same number as the population of *A. lwoffii* sequences) from the *A. baumannii* population 100 times to create an average. *A. baumannii* encodes significantly more ARGs than *A. lwoffii (*p <0.0001); the mean number of ARGs in *A. lwoffii* was 9 but was 30 for *A. baumannii,* Fig. 1a.

**Fig. 1.**
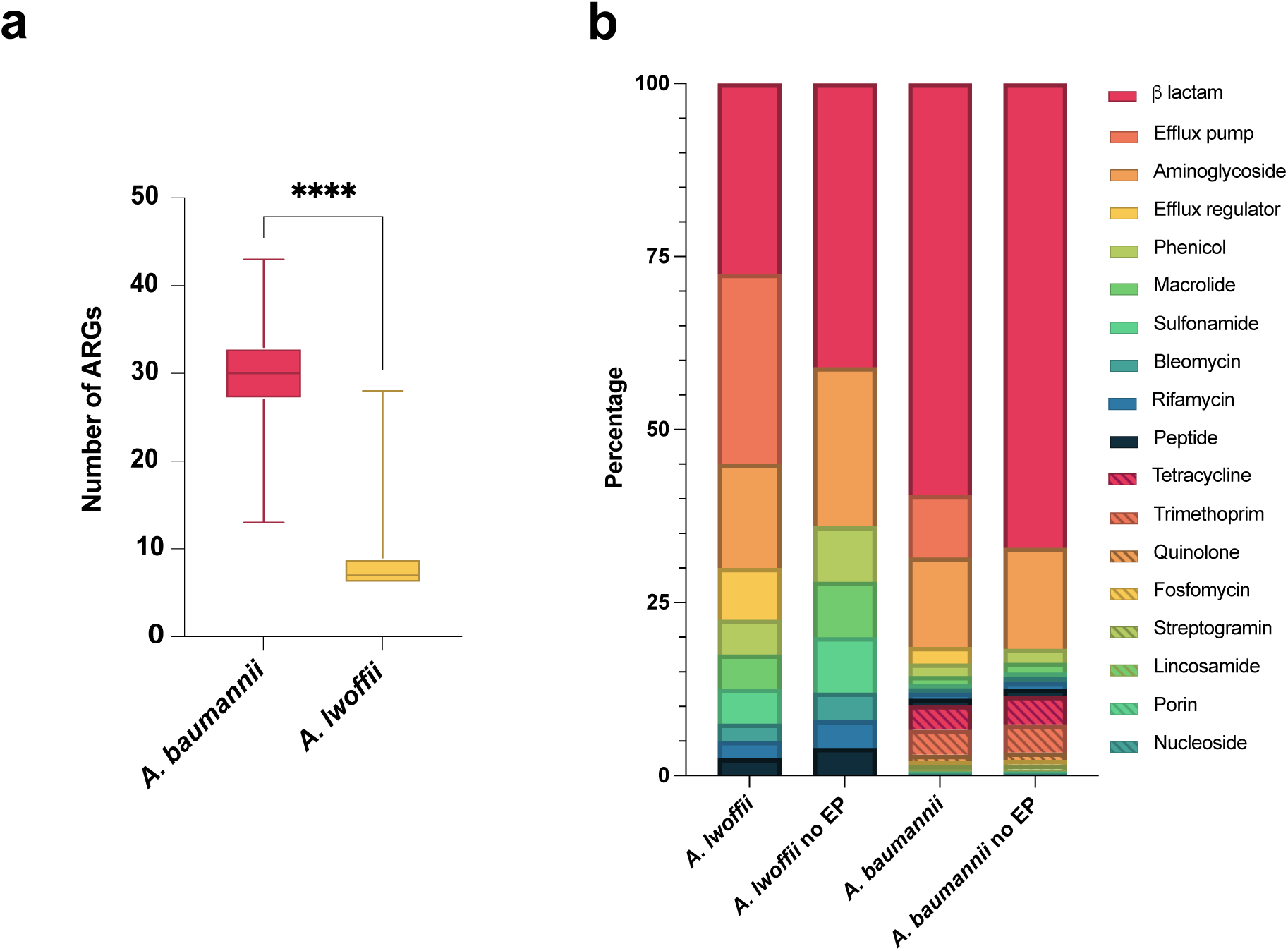
*A. baumannii* encodes more ARGs than *A. lwoffii*. a - Number of ARGs found per whole genome sequence from either species. *A. baumannii* in pink n= 4,809, *A. lwoffii* in yellow n=38. A random permutation and Welch’s T test was performed to compare the average number of genes when the sample sizes were the same - ****, p<0.0001. b – stacked bar chart showing drug classes targeted by all antibiotic resistance genes found in *A. lwoffii* and *A. baumannii* whole genome sequences. Only 40 different ARGs were found for *A. lwoffii* whereas 333 different ARGs were found across *A. baumannii,* but this is likely explained by the different dataset sizes of either species. EP - efflux pump associated genes

Although there was a difference in total gene presence, the classes of antibiotics that the ARGs were active against was similar across the two species, Fig. 1b. The majority of ARGs (>50%) found in *A. lwoffii* and *A. baumannii* reduce the host’s susceptibility to beta lactams and aminoglycosides.

### *A. lwoffii* and *A. baumannii* possess similar genomic signatures of horizontal gene transfer, but *A. lwoffii* contains more DNA defence systems

The greater antibiotic resistance levels of *A. baumannii* are seemingly explained by the fact that this species harbours significantly more ARGs than *A. lwoffii*. However, both species inhabit similar niches, cause similar types of infection, and therefore are expected to have been exposed to similar antibiotics. Variation in rates of horizontal gene transfer into and within each species might explain the difference in the numbers of ARGs they carry. To investigate this, the presence of prophage and plasmid-associated sequences, type IV pili genes for natural transformation and the presence of DNA defence systems, which would limit the acquisition of foreign DNA, were searched for in the whole genome sequences.

To determine whether *A. baumannii* and *A. lwoffii* harbour different numbers or types of plasmids, ABRicate was used to screen for plasmid replicons from an *Acinetobacter* replication initiation (*rep*) gene database (23). An average of 4 and 2 *rep* genes were found per *A. lwoffii* and *A. baumannii* genome, respectively. A random permutation and Welch’s T test revealed that *A. lwoffii* contained significantly more *rep* genes than *A. baumannii* (p <0.0001), suggesting that *A. lwoffii* harbours more plasmids, Fig. 2a.

**Fig. 2.**
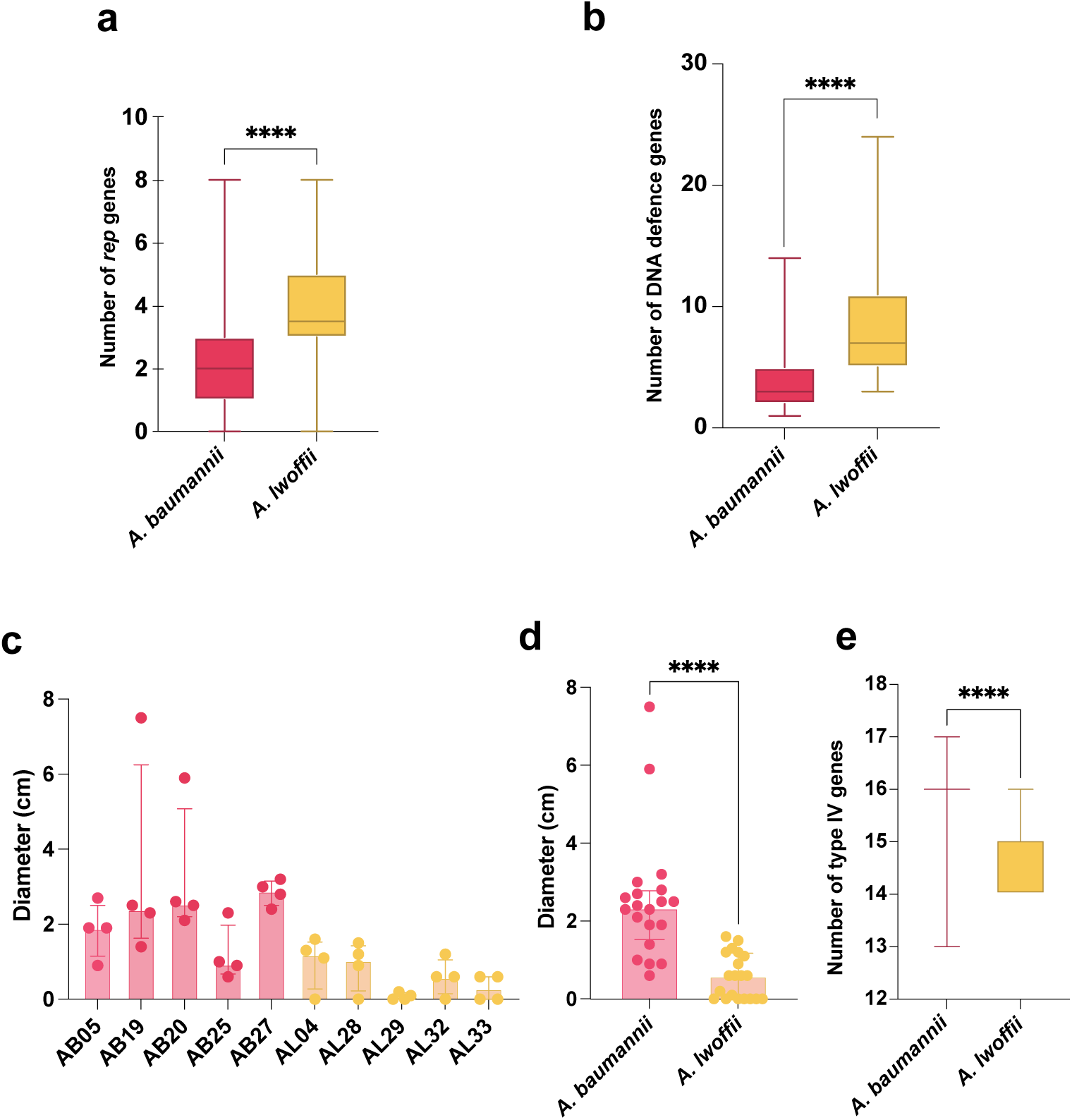
Signatures of foreign DNA acquisition. a - Number of *rep* genes found per whole genome sequence from either species, b – number of DNA defence system genes per sequence. *A. baumannii* in pink n= 4,809,*A. lwoffii* in yellow n=38. c- the twitching motility of individual strains, d- combined values of all strains tested per species, and e- the number of type IV pili associated genes found in the whole genome sequences. A random permutation and Welch’s T-test was used to compare the mean number of *rep,* DNA defence system and type IV associated genes in comparable datasets of *A. baumannii* and *A. lwoffii*, *A. baumannii* genomes encode more genes than *A. lwoffii* (in all instances p <0.0001).

*Acinetobacter rep* genes are classified broadly according to the protein family they encode (Rep_1, Rep_3 or RepPriCT_1) and specifically by homology (>95% nucleotide identity cut-off) to a collection of reference *rep* sequences (23). All *A. lwoffii* rep genes detected here belonged to the Rep_3 (R3) group. However, since the *rep* database was constructed primarily for the purpose of typing plasmids in *A. baumannii,* there were inconsistencies when comparing the *rep* genes identified by ABRicate and the number of circular plasmid sequences in complete *A. lwoffii* genomes. ABRicate detected fewer *rep* genes (n=34) than there were plasmids (n=64) in the complete genomes (supplementary S5). Whilst it is possible that some plasmids did not contain a recognisable *rep* gene, as has been reported for *A. baumannii* plasmids (23), this was unlikely to be the case for all instances here. Therefore, the NCBI annotations for all plasmids in complete *A. lwoffii* genomes were screened for ORFs labelled “rep”, and a further six genes not represented in the database were found, five encoding Rep_3 proteins - CP032104 1 (pALWEK1.11), CP080579 1 (pALWVS1.3), CP072552.1 (pH7-68), CP080580 1 (pALWVS1.4), CP080643 1 (pALWEK1.16) and one encoding Rep_1 (CP080641; pALWEK1.14). In a phylogenetic tree, these genes clustered independently of previously known *rep* genes (supplementary S6). With these considered, all but one *A. lwoffii rep* genes clustered in R3, supporting the idea that *A. lwoffii* almost exclusively maintains R3-type plasmids.

The most common *rep* types in *A. lwoffii* were R3-T25/R3-T45, which were found in a total of 92% of genomes. R3-T25 and R3-T45 are 94.71% identical at the nucleotide level and therefore, although classed as different *rep* types using a 95% cut-off value, are very closely related. Therefore, we propose that R3-T25/R3-T45 replicons represent a native *A. lwoffii* plasmid family, found in almost all complete genome sequences of this species examined here. In contrast, R3-T25/R3-T45 replicons were only found in 0.4% of *A. baumannii* genomes. For *A. baumannii,* 38% of sequences contained R2-T1 and 37% encoded RP-T1 *rep* types. In total, *A. baumannii* had 82 distinct *rep* types, including from RP, R1, R2 and R3 groups. A full list of *rep* genes highlighted in both species’ can be found in supplementary S7.

In addition to ARGs, occasionally, plasmids may also carry genes for RND efflux pumps, which can export a wide range of structurally diverse compounds, including antibiotics (12), and can act as important mechanisms for antibiotic resistance. RND determinants have been seen in plasmids in *A. baumannii,* for example pDETAB2 from a Chinese ICU patient isolate (32), and more recently in *A. lwoffii*, where AL_065, which was isolated from a hospital bed rail in Pakistan, harboured a plasmid encoding an RND transporter and periplasmic adaptor protein (33). This plasmid (CP078046.1, *rep* type R3-T25) is also found in *A. baumannii* and has the potential to disseminate RND efflux genes across *A. lwoffii* more broadly. The RND pump is closest in homology to AdeB (31) and may therefore represent the acquisition of an additional, adaptive RND pump, reducing the susceptibility of this strain to structurally different substrates than those exported by its native RND system: AdeIJK (1).

To determine if the relative lack of ARGs in *A. lwoffii* could also be related to other mechanisms of HGT, we searched for the presence of prophage DNA within genomes of both species. Both *A. lwoffii* and *A. baumannii* had prophage DNA within their genomes, as determined by PHASTER (supplementary S8). Therefore, both species have been previously infected by phage and have the capacity to acquire novel DNA, such as ARGs, introduced by phages.

The number of DNA defence systems across the two species was determined as this could impact their acquisition and maintenance of foreign DNA. Using DefenseFinder *A. lwoffii* genomes were found to encode between 3 and 24 defence systems per genome which was significantly more than *A. baumannii* which had between 1 and 14 (p=<0.0001) (Fig. 2b). The types of defence systems present also differed. *A. lwoffii* encoded mostly type I and IV restriction modification systems, which cleave unmethylated DNA whereas *A. baumannii* encodes more PsyrTA toxin antitoxin systems and antiphage systems e.g. SspBCDE (supplementary S9).

*Acinetobacter* species can display twitching motility in laboratory conditions, which aids the natural transformation of DNA from the extracellular environment into the cell (22). Therefore, the ability of *A. lwoffii* and *A. baumannii* to twitch was measured. While there was strain variation in sub surface twitching motility, generally *A. lwoffii* twitched less (average of 0.6 cm) than *A. baumannii* (average of 2.5 cm) at 37°C, Fig. S2 (c,d), suggesting that *A. lwoffii* may be less naturally competent than *A. baumannii*.

Natural transformation uses type IV pili genes and therefore we also looked for the presence of genes associated with type IV pili in both species (Fig. 2e). There were significantly more type IV associated genes found in *A. baumannii* genomes compared to *A. lwoffii* genomes (p<0.001), supplementary S10.

### *A. baumannii* readily evolved resistance to meropenem, ciprofloxacin and gentamicin but *A. lwoffii* only evolved resistance to ciprofloxacin

Since *A. lwoffii* has remained susceptible to antibiotics, we sought to determine whether it can evolve resistance to clinically relevant antibiotics *in vitro*. For context, we also included *A. baumannii*, which is well known to evolve drug resistance rapidly. To this end, selection experiments were set up, where susceptible strains were grown in the presence of increasing concentrations of meropenem, ciprofloxacin or gentamicin. After 7 days, whole genome sequencing was performed to characterise any genomic changes compared to the ancestral strain (supplementary S11).

*A. baumannii* (AB18) mutants selected in the presence of meropenem had meropenem MICs 2-3-fold above that of the parent strain MIC, from 1 to 2-4 μg/mL (supplementary S11). There were also MIC increases for ampicillin (4-5-fold), ciprofloxacin (3-fold) and tetracycline (3- fold) with some mutants also being less susceptible to moxifloxacin (2-3-fold) and erythromycin (2-3-fold), Fig. 3. It was noted that fewer *A. lwoffii* (AL28) colonies were selected for, however when MIC testing the mutants, the increase was also 3-fold above the ancestral MIC from 0.015 to 0.06 μg/mL. There was no significant MIC change for the other antibiotics tested.

**Fig. 3.**
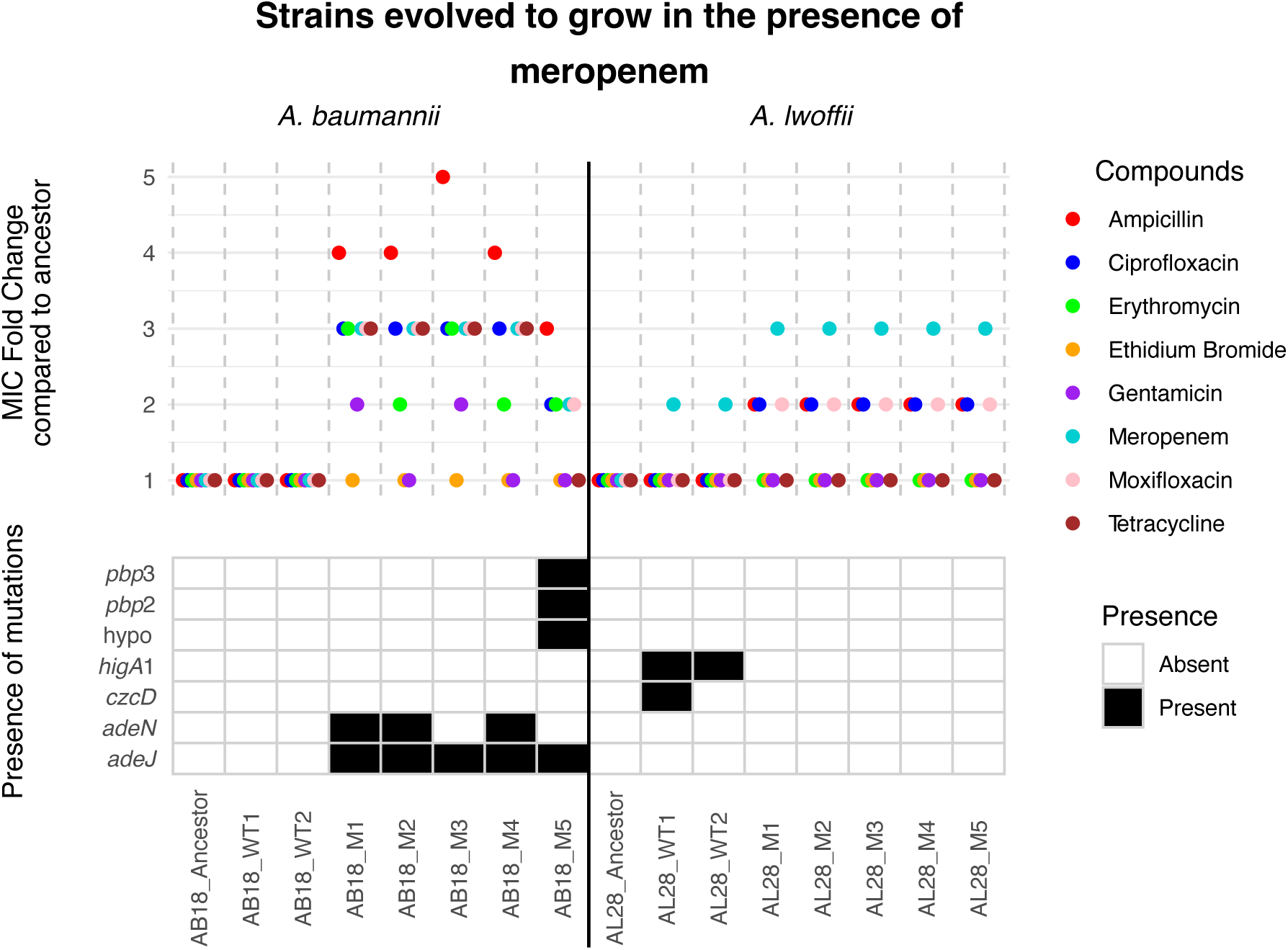
MIC fold change results and SNP presence for strains evolved to grow in increasing concentrations of meropenem. “Ancestor” and “WT” (broth-only) controls are compared to individual “M” (mutant) isolates from the terminal passage. AB18 - *A. baumannii* and AL28 - *A. lwoffii*. An MIC fold change of 1 means the strain is as susceptible or more susceptible to the drug compared with the ancestor.

Five mutants from AL28 and AB18 were subject to whole genome sequencing and their sequences were compared to the ancestral strain and parental strains which had been passaged in the same experiment in nutrient broth only. Despite *A. lwoffii* being able to grow at the final concentration of meropenem used in the evolution experiment no canonical resistance mutations were seen. In fact, no SNPs were found in the mutants, even though the strains passaged in nutrient broth alone encoded some SNPs. However, for *A. baumanii* (AB18) all five sequenced strains had SNPs in the RND efflux transporter encoding gene *adeJ* and in the gene that encodes the repressor protein for this system - *adeN*. Three of the *adeJ* mutations were within the distal binding pocket of the pump, where beta-lactams bind (1). Additionally, AB18 mutant 5 had mutations in genes encoding penicillin binding proteins 2 and 3, known to be involved in meropenem For ciprofloxacin both *A. baumannii* and *A. lwoffii* cultures evolved resistance to above the EUCAST breakpoint. In AB18 large MIC changes, between 9 and 10-fold higher than the ancestral strain, were seen for ciprofloxacin and moxifloxacin. Additionally, MIC increases were also observed for gentamicin (4-5-fold) and erythromycin (3-fold) in some mutants (AB18 M2, M3, M5) and the tetracycline MIC was also elevated (3-fold) in AB18 M2 and M3. Mutants selected in the presence of increasing concentrations of ciprofloxacin had mutations in both the target of the drug (*gyrA/gyrB/parC*) and RND efflux systems (*ade* pumps).

For *A. lwoffii*, in contrast to the results seen with meropenem, target site and efflux SNPs were seen in the AL28 mutants. It is also worth noting the AL28 WT strains also harboured polymorphisms, despite being passaged in nutrient broth only. SNPs were found in genes such as *higA*1, encoding an antitoxin protein, and *yfdX*2, encoding a heat resistance protein. AL28 mutants had SNPs in *adeJ, adeN, atpB, gyrA* and *gyrB*. Presumably, the combination of SNPs in efflux- related genes and target-site genes contributed to the reduced susceptibility of the mutants to ciprofloxacin, moxifloxacin and also protected against meropenem, Fig. 4.

**Fig. 4.**
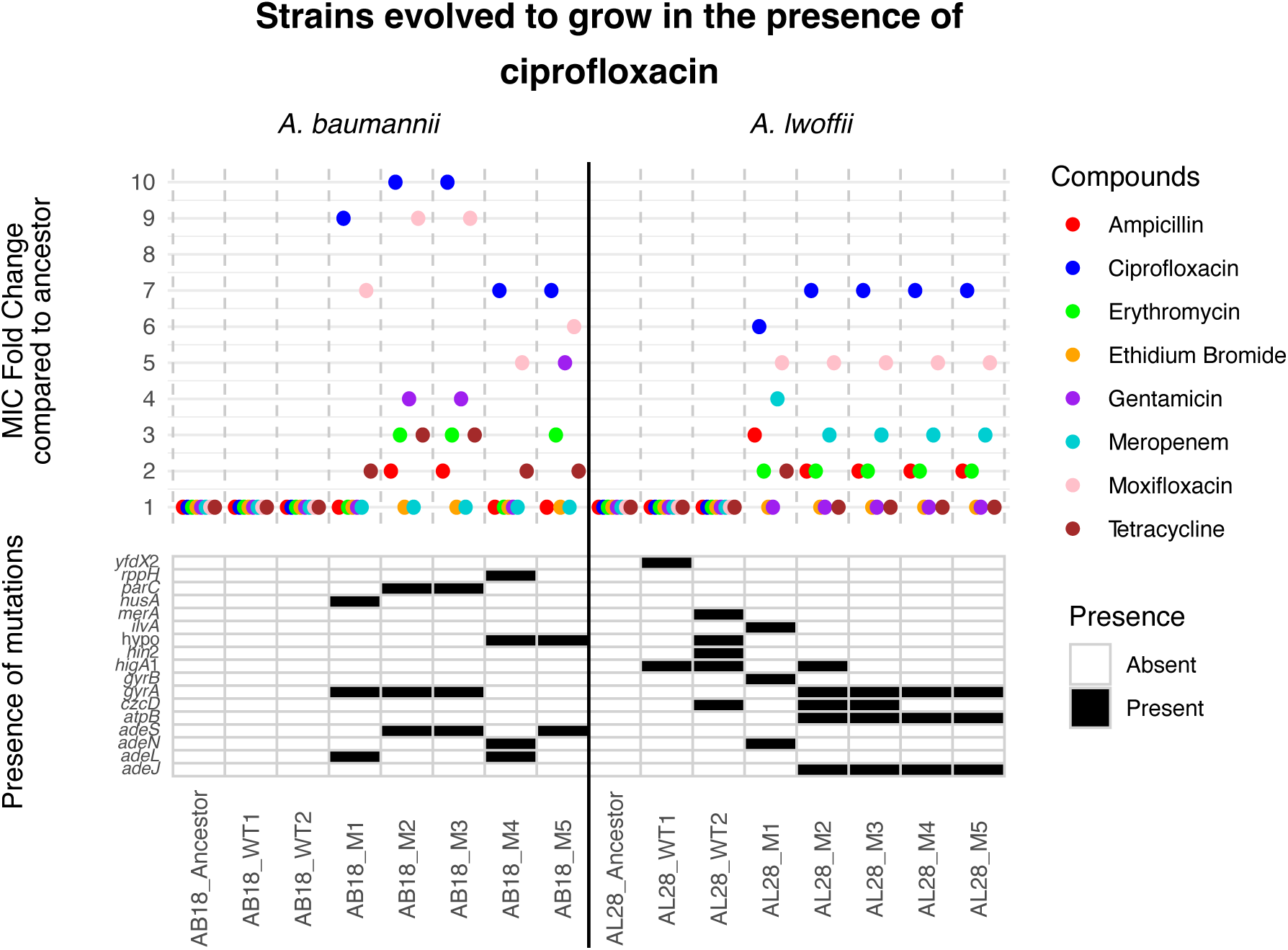
MIC fold change results and SNP presence for strains evolved to grow in increasing concentrations of ciprofloxacin. “Ancestor” and “WT” (broth-only) controls are compared to individual “M” (mutant) isolates from the terminal passage. AB18 - *A. baumannii* and AL28 - *A. lwoffii*. An MIC fold change of 1 means the strain is as susceptible or more susceptible to the drug compared with the ancestor.

Since *A. lwoffii* seemed to be capable of evolving drug resistance mutations to ciprofloxacin but not meropenem a third experiment was conducted. Here, gentamicin was chosen which is also used clinically to treat *Acinetobacter* infections. All AB18 mutants had elevated MICs to gentamicin (8 or 9-fold above ancestral strain MIC), taking them from clinically susceptible to resistant (> 4 μg/mL), Fig. 5. These mutants also displayed a reduced susceptibility to ciprofloxacin and moxifloxacin and some of the AB18 mutants (1, 2 and 4) also showed a reduced susceptibility to erythromycin and tetracycline too. The wild-type strains grown in broth did not encode any SNPs, whereas the mutant strains had SNPs in *adeB, adeR, adeS, fusA, ptsP* and *tetR*.

**Fig. 5.**
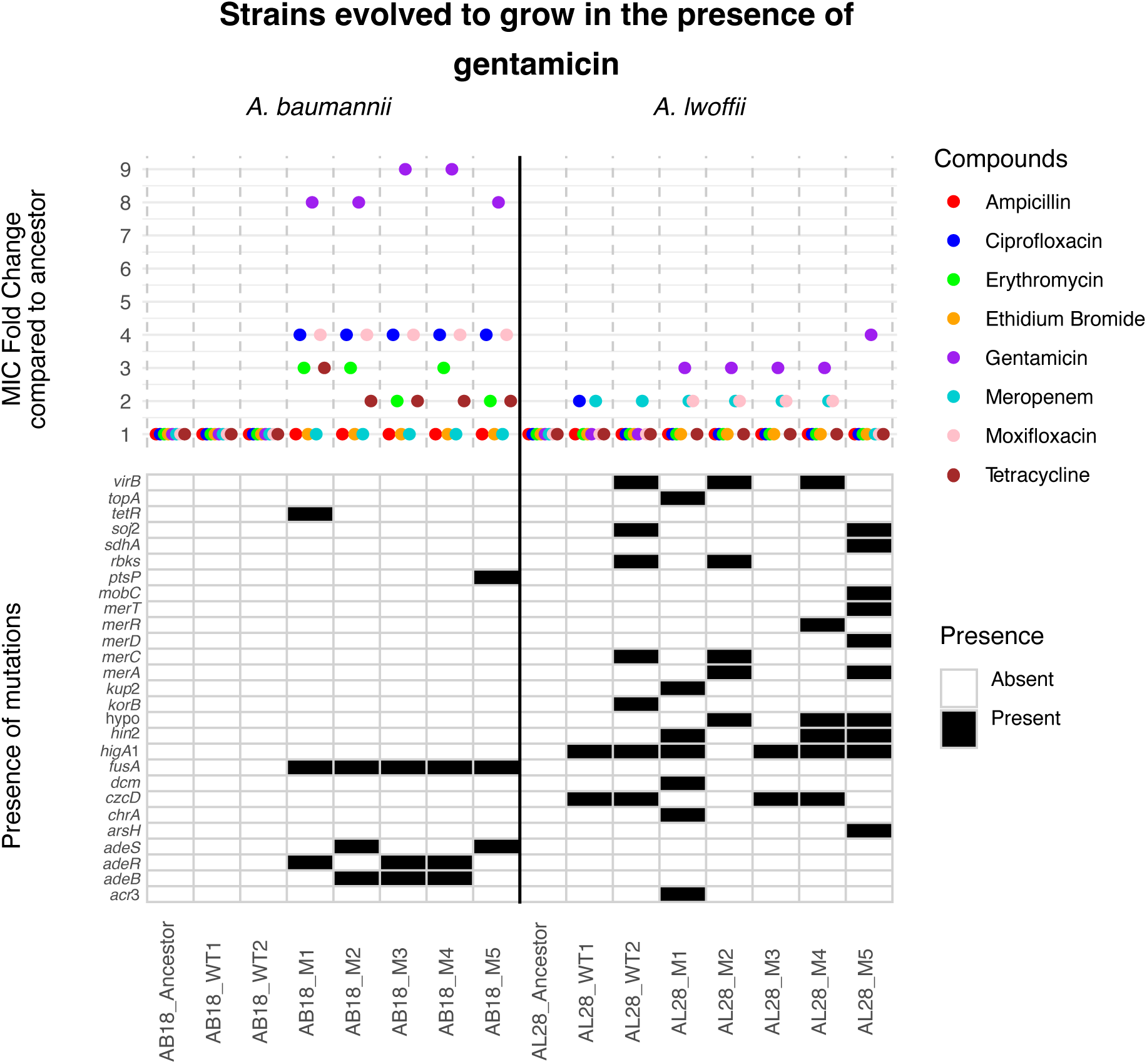
MIC fold change results and SNP presence for strains evolved to grow in increasing concentrations of gentamicin. “Ancestor” and “WT” (broth-only) controls are compared to individual “M” (mutant) isolates from the terminal passage. AB18 - *A. baumannii* and AL28 - *A. lwoffii*. An MIC fold change of 1 means the strain is as susceptible or more susceptible to the drug compared with the ancestor.

As with meropenem, the *A. lwoffii* strain tested did not exhibit drug resistance to gentamicin or other drugs tested. However, during this experiment many SNPs were selected for in both the nutrient broth only conditions (WT1 and WT2) and gentamicin conditions (M1-5). Mutations found only in the AL28 cells grown in gentamicin included SNPs in *acr*3 and *arsH* (arsenic resistance), *chrA* (chromate resistance) and *merA, merD merR* and *merT* (mercuric transport proteins). Therefore, there was both conservative MIC differences and genomic evidence of a stress response, particularly in metal-tolerance genes.

In summary, *A. baumannii* AB18 was able to rapidly evolve resistance to three clinically relevant antibiotics, which provided not only elevated MICs to that antibiotic but also to drugs from other classes. Furthermore, AB18 went from clinically susceptible to resistant, as defined by EUCAST breakpoints, in each instance. However, for *A. lwoffii*, clinical resistance was only seen for ciprofloxacin. These results show that *A. lwoffii* has a more limited capacity to evolve resistance to antibiotics and due to the diversity of efflux-related mutations in *A. baumannii* this may be due to the lack of RND systems in *A. lwoffii*.

### *A. lwoffii* forms less biofilm, and the biofilm is more susceptible to antibiotics and biocides than those formed by *A. baumannii*

Antibiotic susceptibility is known to be decreased when bacteria exist within a biofilm and *Acinetobacter* often forms biofilm to aid survival in the clinical environment (35). Therefore, the biofilm forming capacity and susceptibility of biofilm to antibiotics was determined. In static conditions, *A. baumannii* strains formed significantly more biofilm on average than the *A. lwoffii* strains (p<0.001), median OD_600_ values of 3.39 and 0.53 respectively, supplementary S12 (a, b). However, when biofilm was formed under laminar flow conditions there was no significant difference in the amount of biofilm formed between the two species, supplementary S12c.

When the genomes were searched, for genes previously associated with biofilm formation (24), *A. lwoffii* sequences were found to have a mean of 1 gene per sequence whereas *A. baumannii* had a mean of 8 genes per genome sequence (supplementary S12d, S13). However, as this database was created using genes from *A. baumannii*, biofilm-associated genes exclusive to or uncharacterised in *A. lwoffii* would not have been found using this approach.

Given that a biofilm lifestyle is associated with decreased susceptibility to antibiotics the MIC and minimum biofilm eradication concentration (MBEC) was determined for representatives of both species (Table 2). For both species the MBEC values were generally higher than the MIC values, for example for AB20 the cefotaxime MBEC was 10-fold higher than the MIC. However, the effect was less evident in *A. lwoffii* (AL04), where there were instances where the MBEC and MIC values did not significantly change (chlorhexidine, meropenem and triclosan). Furthermore, in general the *A. lwoffii* (AL04) MBEC values were lower than those of *A. baumannii* (AB20). Therefore, whilst the biofilms formed by both strains were less susceptible to antibiotics and biocides, the biofilm formed by *A. baumannii* (AB20) afforded greater protection than in *A. lwoffii*.

**Table 2.** Minimum biofilm eradication concentrations (MBEC) and minimum broth inhibitory concentrations (MIC) of antibiotics and biocides in *A. lwoffii* (AL04) and *A. baumannii* (AB20).

|  | AL04 |  | AB20 |  |
| --- | --- | --- | --- | --- |
|  | MBEC | MIC | MBEC | MIC |
| Cefotaxime (µg/mL) | 256 | 2 | 8192 | 16 |
| Chlorhexidine (%) | 0.0017 | 0.0008 | <1 | 0.0035 |
| Ciprofloxacin (µg/mL) | 16 | 0.06 | 128 | 2 |
| Meropenem (µg/mL) | 0.06 | 0.03 | <128 | 0.25 |
| Oxacillin (µg/mL) | 8 | 16 | <4096 | 512 |
| Tetracycline (µg/mL) | 256 | 0.5 | 1024 | 2 |
| Triclosan (µg/mL) | 0.5 | 0.5 | <128 | 1 |
| Rifampicin (µg/mL) | 8 | 0.5 | 64 | 4 |

### *A. lwoffii* has a longer cell morphology than *A. baumannii*

Thus far it is clear that *A. lwoffii* is more susceptible to antibiotics than *A. baumannii*, in both static and biofilm conditions and this is likely due to a reduced ability to evolve and acquire resistance, which may be underpinned by the presence of more DNA defence systems and fewer RND efflux pumps. Given the lack of research into *A. lwoffii*, the basic biology of the two species under laboratory conditions was assessed.

To determine whether there were any morphological differences between these two species, two strains of *A. baumannii* (AB05, AB18) and two strains of *A. lwoffii* (AL04, AL28) were imaged using scanning electron microscopy (SEM). *A. lwoffii* had significantly longer cells than *A. baumannii,* (n=100 cell measurements per strain) Fig. 6, supplementary S14.

**Fig. 6.**
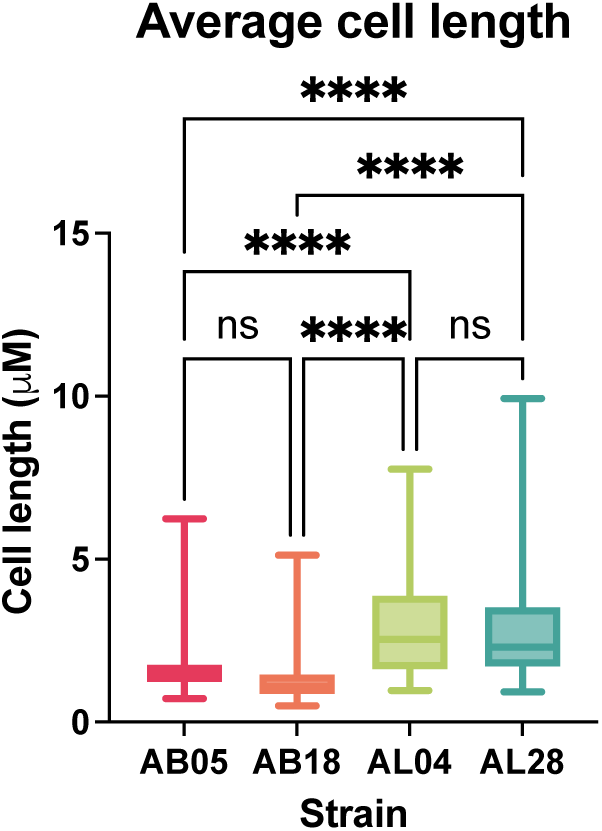
Average cell length (μM) of *A. baumannii* (AB) and *A. lwoffii* (AL) strains imaged by the Apreo 2 Scanning Electron Microscope. *A. lwoffii* had statistically longer cells than *A. baumannii* strains (*****p=<0.0001, one-way ANOVA with Tukey’s multiple comparisons). The whiskers on the box plot show minimum and maximum values obtained.

### *A. baumannii* grows more readily in both LB and human serum than *A. lwoffii*

Additionally, the growth of both species was compared at 37°C, 30°C and 25°C. In LB *A. lwoffii* grew to a lower final density than *A. baumannii* at all temperatures. Growing at cooler temperatures generally increased the length of the lag phase. The mean generation times (supplementary S15) were generally faster at 30°C for *A. lwoffii* while *A. baumannii* grew fastest at 37°C. While *A. lwoffii* grew to a lower final OD than *A. baumannii* (supplementary S16) the generation times of AL28, AL32 and AL33 grew at comparable rates to the *A. baumannii* strains.

Due to the capacity of both species to cause bacteraemia in humans, we also sought to understand how well both species survive and grow in human serum. Growth was compared in human serum with and without complement proteins (normal human serum (NHS) and heat inactivated serum (HIS), respectively); both species grew more slowly in serum than LB (supplementary S17 and S18). Of the two *A. lwoffii* strains tested, AL04 had a prolonged lag but did grow in both HIS and NHS, although growth rate was better in HIS. AL28 did not grow in serum and formed clumps, making OD measurements problematic. *A. baumannii* AB05 and AB18 grew as well in normal serum and as they did in HIS. AB18 grew significantly (p=0.0098) better than AB05 in HIS. All other conditions were not significantly different.

Survival in human serum was also measured, to determine whether, although not actively growing, strains could remain viable in the presence of serum. All strains, except AL28, could survive in NHS and by 24 hours CFU/mL was similar in both serum and the LB control, Fig. 7a.

**Fig. 7.**
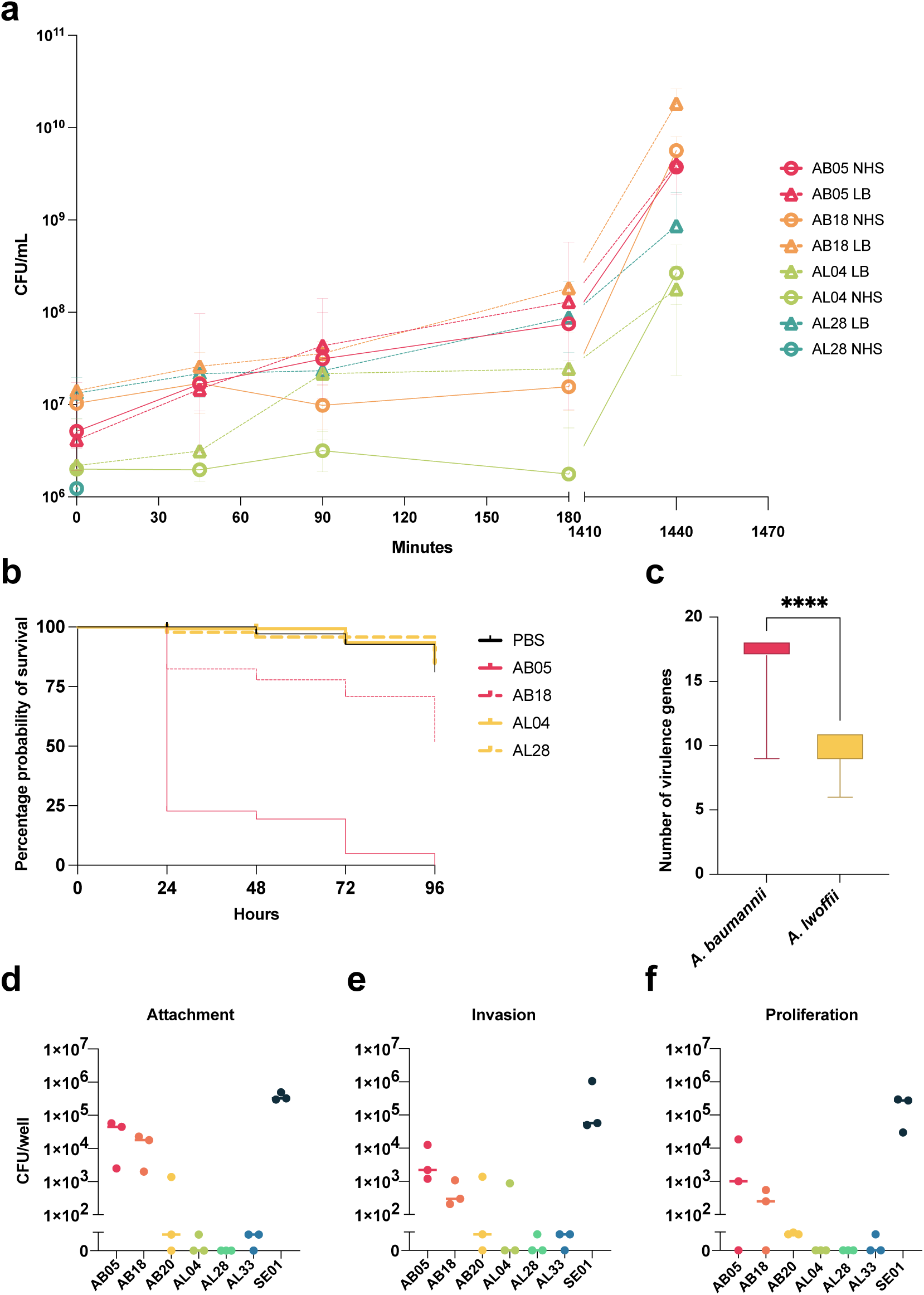
a- the survival of *A. baumannii* (AB) and *A. lwoffii* (AL) strains in both LB (dashed line) and normal human serum (continuous line) over 24 hours. b- Survival of *Galleria mellonella* after inoculation with either *A. baumannii* (AB, pink) or *A. lwoffii* (AL, yellow). A PBS injury control is included in black. c- the number of virulence associated genes found in *A. baumannii* (pink) and *A. lwoffii* (yellow) whole genome sequences. A random permutation and Welch’s T test was used and shows that *A. baumannii* encodes significantly more virulence genes than *A. lwoffii*. Attachment (d), invasion (e) and proliferation (f) of both species in human THP-1 macrophages was measured. SE01 is a positive control of *Salmonella enterica* Typhimurium. AB = *A. baumannii* AL = *A. lwoffii.* Comparative statistics (one-way ANOVA) were performed but no conditions were significantly different.

A synthetic wound model (36) also showed that *A. lwoffii* strains did not grow as well as *A. baumannii* strains. This supports the fact that *A. lwoffii* survived poorly in the presence of human serum as AL04 and AL28 did not grow, supplementary S19.

### A. baumannii is more virulent than A. lwoffii in vivo and in vitro

We also sought to determine whether there was a difference in the *in vivo* virulence capacity of the two species and chose to use the well-characterised infection model organism *Galleria mellonella,* which has an innate immune system (30). When *G. mellonella* larvae were infected with 1x10^6^ *A. baumannii* or *A. lwoffii* cells, more larvae were killed when infected with *A. baumannii* (AB05 and AB18) than *A. lwoffii* (AL04 and AL28) which correlates with what has been seen previously (30). By 48 hours the probability of larvae survival was <25% for AB05 infection, whereas it was >95% for AL28. Of the *A. baumannii* strains, AB05 was significantly more virulent than AB18 in this model, Fig. 7b (p<0.0001, Log-rank test).

Since *A. baumannii* was more virulent *in vivo* than *A. lwoffii*, we also probed the whole genome sequences for the presence of virulence genes. *A. baumannii* genomes encoded significantly more virulence genes than *A. lwoffii* genomes, p<0.0001, (Fig. 7c, supplementary S20) when using a random permutation and T test to compare two equally sized sample sets.

Finally, to determine virulence *in vitro*, strains were incubated with a human macrophage cell line, THP-1. *A. baumannii* strains (AB05 and AB18) were able to attach to and subsequently invade THP-1 cells Fig. 7 (d and e). However, after six hours proliferation was also measured and there was no difference in the number of CFUs between invasion and proliferation suggesting *A. baumannii* was not actively growing within the cells but could survive at least for the period of the assay, Fig. 8f. In contrast, neither of the *A. lwoffii* strains tested could attach to or invade human macrophage cells *in vitro*.

## Discussion

The emerging pathogen, *A. lwoffii* is the leading cause of *Acinetobacter*-derived blood stream infections in England and Wales, followed by the extensively studied *A. baumannii* (4). However, *A. baumannii* has developed widespread multidrug resistance while *A. lwoffii* has remained sensitive to almost all antibiotics. Whilst research into *A. baumannii* is increasing and more is known about its antibiotic resistance there remains a knowledge gap in understanding the emerging opportunistic pathogen *A. lwoffii*. This work aimed to explore differences in the two species in terms of their antibiotic susceptibility, infectivity, and basic biology. We have shown that *A. lwoffii* is more susceptible to drugs used to treat *Acinetobacter* infections than *A. baumannii*, is less virulent and does not evolve drug resistance to the same degree as *A. baumannii*.

This work confirmed previous data suggesting *A. lwoffii* is susceptible to antibiotics while *A. baumannii* is commonly multidrug resistant (4) and showed the difference in phenotype is caused by *A. lwoffii* encoding fewer resistance genes than *A. baumannii*. Both species are found in similar environments such as on the human body, although *A. baumannii* is not considered to be part of a healthy skin microbiome (5, 37). As they are both found within the hospital environment, it is peculiar that resistance (either by mutation or the horizontal acquisition of ARGs) has not been commonly selected for in *A. lwoffii*.

The lack of ARGs in *A. lwoffii* may be due, at least in part, to the presence of DNA defence systems that are absent in *A. baumannii,* such as a greater number of restriction modification systems. The presence of more DNA defence systems in *A. lwoffii* suggests that this species is more stringent about the DNA it maintains (38).

In addition to fewer ARGs, *A. lwoffii* also less readily evolved resistance to three clinically relevant drugs compared to *A. baumannii*. Drug resistance mutations often occur in the drug’s target: penicillin binding proteins for meropenem (34), DNA gyrase for ciprofloxacin (39–41) and the ribosome for gentamicin (42). This was the case for *A. baumannii* here. In the one instance where *A. lwoffii* evolved resistance, to ciprofloxacin, drug target mutations were also observed. Ciprofloxacin mutations often occur in the quinolone resistance determining regions (QRDR) of GyrA, GyrB and ParC (41). The *A. baumannii* mutations in *gyrA* were in the QRDR (amino acids 65-104) but *A. baumannii* mutations in *parC* and *A. lwoffii* in *gyrB,* however, were not within the QRDRs.

Additionally, mutations were captured in RND efflux pumps that export the compounds used for selection. For example, *A. baumannii* meropenem mutants had *adeJ* mutations and beta-lactams bind to the distal pocket of AdeJ (42, 43). Fluoroquinolones can be exported by all three Ade pumps in *A. baumannii* (37), which explains why mutations in all three Ade systems were seen, including mutations that affected the regulators of these systems. Gentamicin is exported by AdeB and can bind to both the proximal and distal binding pockets but Y77, T91, and S134 are thought to be essential for gentamicin binding to the proximal pocket of AdeB (44). Given the proximity of the *A. baumannii* AdeB mutations (amino acids 97 and 136) in this study to those reported in the literature (44), it is likely that these mutations led to increased gentamicin export via AdeB. Mutations in AdeRS have been reported to increase AdeABC expression, for example A91V in AdeR and A94V in AdeS (45). This study also captured the A91V SNP in AdeR, which sits in the signal receiver domain, as well as other mutations in AdeRS, indicating that AdeRS may be being modulated to increase AdeABC expression and the extrusion of gentamicin.

The mutant evolution experiments clearly show that *A. lwoffii* has a reduced capacity to evolve resistance to antibiotics compared to *A. baumannii,* where it only evolved resistance to ciprofloxacin. This could be because *A. lwoffii* only encodes one tripartite RND system (AdeIJK) (1). RND efflux pumps have an underpinning role in the development of resistance via other molecular mechanisms (42). For example, in other species of Gram-negative bacteria deletion of efflux pumps reduces the mutation selection frequency (42, 46). In addition, mutations within efflux pumps often occur first evolutionarily and allow for the development of more canonical drug target mutations, which may have been the case in this study (47, 48). The reduced efflux capacity of *A. lwoffii* could therefore limit the selection of drug resistance mutations. This is further supported by the fact that in *A. baumannii*, drug resistance mutations were found across all three tripartite systems, indicating their important role in resistance evolution. Another potential mechanism for the lack of resistance development could be more stringent DNA repair mechanisms in *A. lwoffii,* for example mismatch repair to inhibit the recombination of non-homologous DNA (49).

When looking at infection-related phenotypes, *A. baumannii* was more virulent than *A. lwoffii*. It was already known that certain *A. baumannii* strains could infect macrophages and persist within their vacuoles, but this was the first time this experiment had been done using *A. lwoffii,* where none of the strains tested could persist within macrophages (31). This could indicate that it is easier to clear *A. lwoffii* infections.

In summary *A. lwoffii* is more susceptible to antibiotics than *A. baumannii* due to a lack of acquired and evolved resistance. Ultimately, an open question remains surrounding why *A. lwoffii* does not seem to be developing drug resistance in the clinic and more work is needed to elucidate if this results from a lack of efflux systems and/or more stringent DNA repair and defence, or other factors. Whilst the widespread antibiotic susceptibility of *A. lwoffii* allows for successful clinical outcomes, there are sporadic cases of drug resistant *A. lwoffii,* highlighting the possibility that drug resistance could emerge (9, 50). It is, therefore, important to fully chart the development of this emerging pathogen to limit the development of drug resistance.

### Author Statements

#### Author contributions

E. M. D. and J.M.A.B. conceptualised and designed the study. E.M.D. carried out genotypic and phenotypic analyses. Bioflux experiments and imaging were done by E. H. Electron microscopy images were taken by T. M with help from L.S. Tissue culture experiments were done with training from B. C and mutant evolution experiments were done with training from R. S. M. Plasmid analysis was provided by R. M. The manuscript was written by E. M. D and J. M. A. B with input from R. M, E. H, B. C., R. S. M, M. A. W and E. M. F.

#### Ethical approval

This study did not require ethical approval.

## Acknowledgements

The authors thank Dr. Rebecca Hall for help with evolution experiments, Dr. Chris Connor for the MASH R script and Dr. Andrew Edwards for help with serum experiments. The authors also thank Dr. Matthew Wand, Dr. Katherine Hardy, Dr. Benjamin Evans and Dr. Jolinda Pollock for kindly sharing *Acinetobacter* strains.

