## Supplementary material for "Differential development of antibiotic resistance and virulence between *Acinetobacter* species": S11

| | Mutation information | | | | | | | | | | | | | MIC Results | | | | | | | |
|---|---|---|---|---|---|---|---|---|---|---|---|---|---|---|---|---|---|---|---|---|---|
| Strain | CHROM | POS | TYPE | REF | ALT | EVIDENCE | FTYPE | STRAND | NT_POS | AA_POS | EFFECT | LOCUS_TAG | PRODUCT | Ampicillin | Ciprofloxacin | Eythromycin | Ethidium Bromide | Gentamicin | Meropenem | Moxifloxacin | Tetracycline |
| AB18 Ancestor | | | | | | | | | | | | | | 16 | 0.25 | 8 | 512 | 0.25 | 1 | 0.06 | 1 |
| AL28 Ancestor | | | | | | | | | | | | | | 0.25 | <0.03 | 0.25 | 8 | 0.06 | 0.015 | <0.015 | 0.25 |
| | HGMGGKAI_16 | 233 | snp | A | G | G:14 A:0 | rRNA | - | | | intergenic_region | HGMGGKAI_03682 | 23S ribosomal RNA (partial) | | | | | | | | |
| Mero_AB18 WT1 | HGMGGKAI_16 | 242 | snp | T | C | C:14 T:0 | rRNA | - | | | intergenic_region | HGMGGKAI_03682 | 23S ribosomal RNA (partial) | 16 | 0.25 | 8 | 512 | 0.25 | 0.25 | 0.06 | 1 |
| Mero_AB18 WT2 | | | | | | | | | | | | | | 16 | 0.25 | 8 | 512 | 0.25 | 0.5 | 0.06 | 1 |
| | HGMGGKAI_1 | 416838 | del | TA | T | T:18 TA:0 | CDS | - | 362/654 | 121/217 | frameshift_variant c.362delT p.Leu121fs | HGMGGKAI_00408 | AdeN | | | | | | | | |
| Mero_AB18 M1 | HGMGGKAI_5 | 211376 | snp | A | G | G:20 A:0 | CDS | - | 416/3177 | 139/1058 | missense_variant c.416T>C p.Val139Ala | HGMGGKAI_02852 | AdeJ | 128 | 1 | 32 | 512 | 0.5 | 4 | 0.25 | 4 |
| | HGMGGKAI_1 | 416914 | del | GA | G | G:20 GA:0 | CDS | - | 286/654 | 96/217 | frameshift_variant c.286delT p.Ser96fs | HGMGGKAI_00408 | AdeN | | | | | | | | |
| Mero_AB18 M2 | HGMGGKAI_5 | 210956 | snp | G | A | A:20 G:0 | CDS | - | 836/3177 | 279/1058 | missense_variant c.836C>T p.Ser279Leu | HGMGGKAI_02852 | AdeJ | 128 | 1 | 16 | 512 | 0.25 | 4 | 0.25 | 4 |
| Mero_AB18 M3 | HGMGGKAI_5 | 211377 | snp | C | A | A:20 C:0 | CDS | - | 415/3177 | 139/1058 | missense_variant c.415G>T p.Val139Phe | HGMGGKAI_02852 | AdeJ | 256 | 1 | 32 | 512 | 0.5 | 4 | 0.25 | 4 |
| | HGMGGKAI_1 | 416672 | snp | C | T | T:20 C:0 | CDS | - | 529/654 | 177/217 | missense_variant c.529G>A p.Gly177Arg | HGMGGKAI_00408 | AdeN | | | | | | | | |
| | HGMGGKAI_5 | 211320 | snp | C | A | A:20 C:0 | CDS | - | 472/3177 | 158/1058 | missense_variant c.472G>T p.Val158Leu | HGMGGKAI_02852 | AdeJ | | | | | | | | |
| Mero_AB18 M4 | HGMGGKAI_366 | 387 | snp | G | A | A:12 G:0 | | | | | | | | 128 | 1 | 16 | 512 | 0.25 | 4 | 0.25 | 4 |
| | HGMGGKAI_2 | 309478 | snp | G | C | C:20 G:0 | CDS | + | 1693/1833 | 565/610 | missense_variant c.1693G>C p.Val565Leu | HGMGGKAI_01775 | Peptidoglycan D,D-transpeptidase PBP3 | | | | | | | | |
| | HGMGGKAI_5 | 211386 | snp | A | C | C:20 A:0 | CDS | - | 406/3177 | 136/1058 | missense_variant c.406T>G p.Phe136Val | HGMGGKAI_02852 | AdeJ | | | | | | | | |
| | HGMGGKAI_5 | 256181 | snp | A | G | G:14 A:0 | CDS | - | 44/264 | 15/87 | missense_variant c.44T>C p.Ile15Thr | HGMGGKAI_02892 | hypothetical protein | | | | | | | | |
| Mero_AB18 M5 | HGMGGKAI_6 | 15877 | del | TTA | T | T:18 TTA:0 | CDS | + | 122/2019 | 41/672 | frameshift_variant c.122_123delAT p.Tyr41fs | HGMGGKAI_02904 | Peptidoglycan D,D-transpeptidase PBP2 | 64 | 0.5 | 16 | 512 | 0.25 | 2 | 0.12 | 1 |
| | JPHHDCAP_1 | 62161 | snp | T | C | C:20 T:0 | | | | | | | | | | | | | | | |
| | JPHHDCAP_3 | 26899 | snp | C | A | A:15 C:0 | | | | | | | | | | | | | | | |
| | JPHHDCAP_3 | 26943 | snp | G | C | C:12 G:0 | | | | | | | | | | | | | | | |
| | JPHHDCAP_3 | 41687 | snp | T | C | C:26 T:0 | | | | | | | | | | | | | | | |
| | JPHHDCAP_3 | 42008 | snp | A | T | T:20 A:0 | CDS | + | 72/303 | 24/100 | synonymous_variant c.72A>T p.Ala24Ala | JPHHDCAP_03198 | HigA1 | | | | | | | | |
| Mero_AL28 WT1 | JPHHDCAP_4 | 76996 | snp | C | T | T:12 C:0 | CDS | - | 238/1281 | 80/426 | missense_variant c.238G>A p.Asp80Asn | JPHHDCAP_03293 | CzcD | 0.25 | <0.03 | 0.25 | 8 | 0.06 | 0.03 | <0.015 | 0.25 |
| | JPHHDCAP_3 | 41687 | snp | T | C | C:16 T:0 | | | | | | | | | | | | | | | |
| Mero_AL28 WT2 | JPHHDCAP_3 | 42008 | snp | A | T | T:20 A:0 | CDS | + | 72/303 | 24/100 | synonymous_variant c.72A>T p.Ala24Ala | JPHHDCAP_03198 | HigA1 | 0.25 | <0.03 | 0.25 | 8 | 0.06 | 0.03 | <0.015 | 0.25 |
| Mero_AL28 M1 | | | | | | | | | | | | | | 0.5 | 0.06 | 0.25 | 8 | 0.06 | 0.06 | 0.03 | <0.12 |
| Mero_AL28 M2 | JPHHDCAP_2 | 1976693 | del | CT | C | C:19 CT:0 | CDS | - | 46/1359 | 16/452 | frameshift_variant c.46delA p.Arg16fs | JPHHDCAP_01997 | PhoR | 0.5 | 0.06 | 0.25 | 8 | 0.06 | 0.06 | 0.03 | <0.12 |
| Mero_AL28 M3 | | | | | | | | | | | | | | 0.5 | 0.06 | 0.25 | 8 | 0.06 | 0.06 | 0.03 | <0.12 |
| Mero_AL28 M4 | JPHHDCAP_1 | 62161 | snp | T | C | C:20 T:0 | | | | | | | | 0.5 | 0.06 | 0.25 | 8 | 0.06 | 0.06 | 0.03 | <0.12 |
| Mero_AL28 M5 | | | | | | | | | | | | | | 0.5 | 0.06 | 0.25 | 8 | 0.06 | 0.06 | 0.03 | <0.12 |
| Cipro_AB18 WT1 | | | | | | | | | | | | | | 16 | 0.25 | 8 | 512 | 0.25 | 1 | 0.06 | 1 |
| Cipro_AB18 WT2 | | | | | | | | | | | | | | 16 | 0.25 | 8 | 512 | 0.25 | 1 | 0.06 | 1 |
| | HGMGGKAI_1 | 11670 | snp | G | T | T:20 G:0 | CDS | + | 833/1032 | 278/343 | missense_variant c.833G>T p.Arg278Leu | HGMGGKAI_00009 | AdeL | | | | | | | | |
| | HGMGGKAI_3 | 314460 | snp | C | A | A:20 C:0 | CDS | - | 235/2715 | 79/904 | missense_variant c.235G>T p.Gly79Cys | HGMGGKAI_02241 | GyrA | | | | | | | | |
| Cipro_AB18 M1 | HGMGGKAI_4 | 305711 | del | CAATAAGCATTTCAGGCTT | C | C:18 CAATAA | CDS | - | 633/1485 | 206/494 | conservative_inframe_deletion c.616_633delAAGCCTGAAATGCTTATT p.Lys2( | HGMGGKAI_02597 | NusA | 16 | 64 | 8 | 512 | 0.12 | 0.25 | 4 | 2 |
| | HGMGGKAI_1 | 671516 | snp | C | G | G:20 C:0 | CDS | - | 88/1074 | 30/357 | missense_variant c.88G>C p.Gly30Arg | HGMGGKAI_00655 | AdeS | | | | | | | | |
| | HGMGGKAI_3 | 314453 | snp | G | A | A:20 G:0 | CDS | - | 242/2715 | 81/904 | missense_variant c.242C>T p.Ser81Leu | HGMGGKAI_02241 | GyrA | | | | | | | | |
| Cipro_AB18 M2 | HGMGGKAI_9 | 19938 | snp | T | A | A:20 T:0 | CDS | + | 996/2220 | 332/739 | missense_variant c.996T>A p.Asn332Lys | HGMGGKAI_03384 | ParC | 32 | 128 | 32 | 512 | 2 | 1 | 16 | 4 |
| | HGMGGKAI_1 | 671516 | snp | C | G | G:20 C:0 | CDS | - | 88/1074 | 30/357 | missense_variant c.88G>C p.Gly30Arg | HGMGGKAI_00655 | AdeS | | | | | | | | |
| | HGMGGKAI_3 | 314453 | snp | G | A | A:20 G:0 | CDS | - | 242/2715 | 81/904 | missense_variant c.242C>T p.Ser81Leu | HGMGGKAI_02241 | GyrA | | | | | | | | |
| Cipro_AB18 M3 | HGMGGKAI_9 | 19938 | snp | T | A | A:20 T:0 | CDS | + | 996/2220 | 332/739 | missense_variant c.996T>A p.Asn332Lys | HGMGGKAI_03384 | ParC | 32 | 128 | 32 | 512 | 2 | 1 | 16 | 4 |
| | HGMGGKAI_1 | 11720 | snp | T | A | A:20 T:0 | CDS | + | 883/1032 | 295/343 | missense_variant c.883T>A p.Phe295Ile | HGMGGKAI_00009 | AdeL | | | | | | | | |
| | HGMGGKAI_1 | 416694 | del | ATCAGGAGATG | A | A:18 ATCAG( | CDS | - | 506/654 | 166/217 | frameshift_variant c.497_506delCATCTCCTGA p.Thr166fs | HGMGGKAI_00408 | AdeN | | | | | | | | |
| | HGMGGKAI_1 | 1080609 | snp | G | A | A:10 G:0 | CDS | - | 768/1236 | 256/411 | synonymous_variant c.768C>T p.Gly256Gly | HGMGGKAI_01047 | hypothetical protein | | | | | | | | |
| | HGMGGKAI_4 | 29576 | snp | A | G | G:20 A:0 | | | | | | | | | | | | | | | |
| Cipro_AB18 M4 | HGMGGKAI_4 | 215409 | snp | G | A | A:20 G:0 | CDS | + | 377/486 | 126/161 | stop_gained c.377G>A p.Trp126* | HGMGGKAI_02523 | RppH | 16 | 16 | 8 | 512 | 0.12 | 0.5 | 1 | 2 |
| | HGMGGKAI_1 | 671171 | del | CATTCCAAACCTG | C | C:18 CATTCC | CDS | - | 432/1074 | 141/357 | conservative_inframe_deletion c.421_432delCAGGTTTGGAAT p.Gln141_Asn | HGMGGKAI_00655 | AdeS | | | | | | | | |
| | HGMGGKAI_1 | 1080636 | snp | G | T | T:20 G:0 | CDS | - | 741/1236 | 247/411 | missense_variant c.741C>A p.Asp247Glu | HGMGGKAI_01047 | hypothetical protein | | | | | | | | |
| | HGMGGKAI_1 | 1216275 | snp | G | T | T:20 G:0 | CDS | - | 41/1128 | 14/375 | missense_variant c.41C>A p.Ala14Asp | HGMGGKAI_01168 | hypothetical protein | | | | | | | | |
| | HGMGGKAI_2 | 443324 | snp | T | C | C:20 T:0 | | | | | | | | | | | | | | | |
| | HGMGGKAI_2 | 443328 | ins | T | TCA | TCA:18 T:0 | | | | | | | | | | | | | | | |
| Cipro_AB18 M5 | HGMGGKAI_22 | 1525 | snp | G | A | A:14 G:0 | rRNA | - | | | intergenic_region | HGMGGKAI_03697 | 23S ribosomal RNA (partial) | 8 | 16 | 32 | 512 | 4 | 1 | 2 | 2 |
| | JPHHDCAP_1 | 65985 | snp | C | A | A:20 C:0 | CDS | + | 502/888 | 168/295 | missense_variant c.502C>A p.Pro168Thr | JPHHDCAP_00066 | YfdX2 | | | | | | | | |
| | JPHHDCAP_1 | 65990 | snp | A | T | T:20 A:0 | CDS | + | 507/888 | 169/295 | synonymous_variant c.507A>T p.Ala169Ala | JPHHDCAP_00066 | YfdX2 | | | | | | | | |
| | JPHHDCAP_2 | 2669688 | snp | T | A | A:10 T:0 | | | | | | | | | | | | | | | |
| | JPHHDCAP_3 | 41687 | snp | T | C | C:21 T:0 | | | | | | | | | | | | | | | |
| Cipro_AL28 WT1 | JPHHDCAP_3 | 42008 | snp | A | T | T:21 A:0 | CDS | + | 72/303 | 24/100 | synonymous_variant c.72A>T p.Ala24Ala | JPHHDCAP_03198 | HigA1 | 0.12 | <0.03 | 0.25 | 8 | 0.06 | 0.015 | <0.015 | <0.12 |
| | JPHHDCAP_2 | 1859296 | snp | G | A | A:12 G:0 | | | | | | | | | | | | | | | |
| | JPHHDCAP_3 | 29282 | snp | T | G | G:20 T:0 | CDS | + | 204/1686 | 68/561 | missense_variant c.204T>G p.Asp68Glu | JPHHDCAP_03182 | MerA | | | | | | | | |
| | JPHHDCAP_3 | 30025 | snp | T | G | G:20 T:0 | CDS | + | 947/1686 | 316/561 | missense_variant c.947T>G p.Val316Gly | JPHHDCAP_03182 | MerA | | | | | | | | |
| | JPHHDCAP_3 | 31473 | ins | A | AGTT | AGTT:18 A:0 | CDS | + | 33/213 | Nov-70 | disruptive_inframe_insertion c.30_32dupTTG p.Cys11dup | JPHHDCAP_03185 | hypothetical protein | | | | | | | | |
| | JPHHDCAP_3 | 37827 | snp | A | G | G:22 A:0 | CDS | + | 514/600 | 172/199 | missense_variant c.514A>G p.Arg172Gly | JPHHDCAP_03193 | Hin2 | | | | | | | | |
| | JPHHDCAP_3 | 41687 | snp | T | C | C:26 T:0 | | | | | | | | | | | | | | | |
| | JPHHDCAP_3 | 42008 | snp | A | T | T:20 A:0 | CDS | + | 72/303 | 24/100 | synonymous_variant c.72A>T p.Ala24Ala | JPHHDCAP_03198 | HigA1 | | | | | | | | |
| Cipro_Al28 WT2 | JPHHDCAP_4 | 76996 | snp | C | T | T:12 C:0 | CDS | - | 238/1281 | 80/426 | missense_variant c.238G>A p.Asp80Asn | JPHHDCAP_03293 | CzcD | 0.12 | <0.03 | 0.25 | 8 | 0.06 | 0.015 | <0.015 | <0.12 |
| | JPHHDCAP_2 | 387064 | ins | T | TGA | TGA:20 T:0 | CDS | + | 138/648 | 46/215 | frameshift_variant c.136_137dupGA p.Asp46fs | JPHHDCAP_00481 | AdeN | | | | | | | | |
| | JPHHDCAP_2 | 397197 | del | AATATTACTGAATTTAACT | A | A:16 AATATT | CDS | + | 1083/1539 | 361/512 | disruptive_inframe_deletion c.1083_1100delTATTACTGAATTTAACTA p.Ile362 | JPHHDCAP_00491 | IlvA | | | | | | | | |
| Cipro_AL28 M1 | JPHHDCAP_2 | 1834404 | snp | C | T | T:20 C:0 | CDS | + | 1354/2472 | 452/823 | missense_variant c.1354C>T p.Arg452Cys | JPHHDCAP_01865 | GyrB | 1 | 1 | 0.5 | 8 | 0.03 | 0.12 | 0.25 | 0.5 |
| | JPHHDCAP_2 | 1994600 | snp | C | T | T:20 C:0 | CDS | + | 680/876 | 227/291 | missense_variant c.680C>T p.Ser227Leu | JPHHDCAP_02013 | AtpB | | | | | | | | |
| | JPHHDCAP_2 | 2593005 | snp | C | T | T:20 C:0 | CDS | - | 383/3177 | 128/1058 | missense_variant c.383G>A p.Arg128Gln | JPHHDCAP_02605 | AdeJ | | | | | | | | |
| | JPHHDCAP_2 | 2702012 | snp | C | A | A:20 C:0 | CDS | + | 242/2982 | 81/993 | missense_variant c.242C>A p.Ser81Tyr | JPHHDCAP_02719 | GyrA | | | | | | | | |
| | JPHHDCAP_3 | 41687 | snp | T | C | C:26 T:0 | | | | | | | | | | | | | | | |
| | JPHHDCAP_3 | 42008 | snp | A | T | T:20 A:0 | CDS | + | 72/303 | 24/100 | synonymous_variant c.72A>T p.Ala24Ala | JPHHDCAP_03198 | HigA1 | | | | | | | | |
| Cipro_AL28 M2 | JPHHDCAP_4 | 76996 | snp | C | T | T:12 C:0 | CDS | - | 238/1281 | 80/426 | missense_variant c.238G>A p.Asp80Asn | JPHHDCAP_03293 | CzcD | 0.5 | 2 | 0.5 | 8 | 0.03 | 0.06 | 0.25 | 0.25 |
| | JPHHDCAP_2 | 1994600 | snp | C | T | T:20 C:0 | CDS | + | 680/876 | 227/291 | missense_variant c.680C>T p.Ser227Leu | JPHHDCAP_02013 | AtpB | | | | | | | | |
| | JPHHDCAP_2 | 2593005 | snp | C | T | T:20 C:0 | CDS | - | 383/3177 | 128/1058 | missense_variant c.383G>A p.Arg128Gln | JPHHDCAP_02605 | AdeJ | | | | | | | | |
| | JPHHDCAP_2 | 2702012 | snp | C | A | A:20 C:0 | CDS | + | 242/2982 | 81/993 | missense_variant c.242C>A p.Ser81Tyr | JPHHDCAP_02719 | GyrA | | | | | | | | |
| | JPHHDCAP_2 | 2863918 | ins | A | AT | AT:18 A:0 | | | | | | | | | | | | | | | |
| Cipro_AL28 M3 | JPHHDCAP_4 | 76996 | snp | C | T | T:12 C:0 | CDS | - | 238/1281 | 80/426 | missense_variant c.238G>A p.Asp80Asn | JPHHDCAP_03293 | CzcD | 0.5 | 2 | 0.5 | 8 | 0.03 | 0.06 | 0.25 | 0.25 |
| | JPHHDCAP_2 | 1994600 | snp | C | T | T:20 C:0 | CDS | + | 680/876 | 227/291 | missense_variant c.680C>T p.Ser227Leu | JPHHDCAP_02013 | AtpB | | | | | | | | |
| | JPHHDCAP_2 | 2593005 | snp | C | T | T:20 C:0 | CDS | - | 383/3177 | 128/1058 | missense_variant c.383G>A p.Arg128Gln | JPHHDCAP_02605 | AdeJ | | | | | | | | |
| Cipro_AL28 M4 | JPHHDCAP_2 | 2702012 | snp | C | A | A:20 C:0 | CDS | + | 242/2982 | 81/993 | missense_variant c.242C>A p.Ser81Tyr | JPHHDCAP_02719 | GyrA | 0.5 | 2 | 0.5 | 8 | 0.03 | 0.06 | 0.25 | 0.25 |
| | JPHHDCAP_2 | 1994600 | snp | C | T | T:20 C:0 | CDS | + | 680/876 | 227/291 | missense_variant c.680C>T p.Ser227Leu | JPHHDCAP_02013 | AtpB | | | | | | | | |
| | JPHHDCAP_2 | 2593005 | snp | C | T | T:20 C:0 | CDS | - | 383/3177 | 128/1058 | missense_variant c.383G>A p.Arg128Gln | JPHHDCAP_02605 | AdeJ | | | | | | | | |
| Cipro_AL28 M5 | JPHHDCAP_2 | 2702012 | snp | C | A | A:20 C:0 | CDS | + | 242/2982 | 81/993 | missense_variant c.242C>A p.Ser81Tyr | JPHHDCAP_02719 | GyrA | 0.5 | 2 | 0.5 | 8 | 0.03 | 0.06 | 0.25 | 0.25 |
| Gen_AB18 WT1 | | | | | | | | | | | | | | 16 | 0.25 | 8 | 512 | 0.25 | 0.5 | 0.06 | 1 |
| Gen_AB18 WT2 | HGMGGKAI_1 | 1442472 | snp | C | A | A:14 C:0 | | | | | | | | 16 | 0.25 | 8 | 512 | 0.25 | 0.25 | 0.06 | 1 |
| | HGMGGKAI_1 | 307851 | snp | C | T | T:20 C:0 | CDS | - | 378/579 | 126/192 | stop_gained c.378G>A p.Trp126* | HGMGGKAI_00283 | TetR | | | | | | | | |
| | HGMGGKAI_1 | 672102 | snp | C | T | T:20 C:0 | CDS | - | 277/744 | 93/247 | missense_variant c.277G>A p.Asp93Asn | HGMGGKAI_00656 | AdeR | | | | | | | | |
| | HGMGGKAI_7 | 806 | snp | C | T | T:20 C:0 | CDS | - | 1543/2139 | 515/712 | missense_variant c.1543G>A p.Gly515Ser | HGMGGKAI_03070 | FusA | | | | | | | | |
| Gen_AB18 M1 | HGMGGKAI_366 | 246 | snp | A | T | T:24 A:0 | | | | | | | | 8 | 2 | 32 | 512 | 32 | 1 | 0.5 | 4 |
| | HGMGGKAI_1 | 671425 | snp | T | C | C:20 T:0 | CDS | - | 179/1074 | 60/357 | missense_variant c.179A>G p.Asp60Gly | HGMGGKAI_00655 | AdeS | | | | | | | | |
| | HGMGGKAI_1 | 674000 | snp | C | T | T:20 C:0 | CDS | + | 290/3111 | 97/1036 | missense_variant c.290C>T p.Pro97Leu | HGMGGKAI_00658 | AdeB | | | | | | | | |
| Gen_AB18 M2 | HGMGGKAI_7 | 703 | snp | A | G | G:20 A:0 | CDS | - | 1646/2139 | 549/712 | missense_variant c.1646T>C p.Phe549Ser | HGMGGKAI_03070 | FusA | 16 | 2 | 32 | 512 | 32 | 1 | 0.5 | 2 |
| | HGMGGKAI_1 | 672107 | snp | G | A | A:20 G:0 | CDS | - | 272/744 | 91/247 | missense_variant c.272C>T p.Ala91Val | HGMGGKAI_00656 | AdeR | | | | | | | | |
| | HGMGGKAI_1 | 674117 | snp | G | A | A:20 G:0 | CDS | + | 407/3111 | 136/1036 | missense_variant c.407G>A p.Gly136Glu | HGMGGKAI_00658 | AdeB | | | | | | | | |
| Gen_AB18 M3 | HGMGGKAI_7 | 746 | snp | A | T | T:20 A:0 | CDS | - | 1603/2139 | 535/712 | missense_variant c.1603T>A p.Phe535Ile | HGMGGKAI_03070 | FusA | 16 | 2 | 16 | 256 | 64 | 1 | 0.5 | 2 |
| | HGMGGKAI_1 | 672107 | snp | G | A | A:20 G:0 | CDS | - | 272/744 | 91/247 | missense_variant c.272C>T p.Ala91Val | HGMGGKAI_00656 | AdeR | | | | | | | | |
| | HGMGGKAI_1 | 674117 | snp | G | A | A:20 G:0 | CDS | + | 407/3111 | 136/1036 | missense_variant c.407G>A p.Gly136Glu | HGMGGKAI_00658 | AdeB | | | | | | | | |
| Gen_AB18 M4 | HGMGGKAI_7 | 746 | snp | A | T | T:20 A:0 | CDS | - | 1603/2139 | 535/712 | missense_variant c.1603T>A p.Phe535Ile | HGMGGKAI_03070 | FusA | 16 | 2 | 32 | 256 | 64 | 1 | 0.5 | 2 |
| | HGMGGKAI_1 | 671161 | snp | G | A | A:20 G:0 | CDS | - | 443/1074 | 148/357 | missense_variant c.443C>T p.Ala148Val | HGMGGKAI_00655 | AdeS | | | | | | | | |
| | HGMGGKAI_4 | 215672 | snp | G | T | T:20 G:0 | CDS | + | 70/2295 | 24/764 | stop_gained c.70G>T p.Glu24* | HGMGGKAI_02524 | PtsP | | | | | | | | |
| Gen_AB18 M5 | HGMGGKAI_7 | 806 | snp | C | T | T:20 C:0 | CDS | - | 1543/2139 | 515/712 | missense_variant c.1543G>A p.Gly515Ser | HGMGGKAI_03070 | FusA | 16 | 2 | 16 | 512 | 32 | 1 | 0.5 | 2 |
| | JPHHDCAP_2 | 557129 | snp | T | G | G:12 T:0 | | | | | | | | | | | | | | | |
| | JPHHDCAP_3 | 26899 | snp | C | A | A:14 C:0 | | | | | | | | | | | | | | | |
| | JPHHDCAP_3 | 30707 | snp | G | T | T:20 G:0 | CDS | + | 1629/1686 | 543/561 | missense_variant c.1629G>T p.Lys543Asn | JPHHDCAP_03182 | MerA | | | | | | | | |
| | JPHHDCAP_3 | 37827 | snp | A | G | G:22 A:0 | CDS | + | 514/600 | 172/199 | missense_variant c.514A>G p.Arg172Gly | JPHHDCAP_03193 | Hin2 | | | | | | | | |
| | JPHHDCAP_3 | 41687 | snp | T | C | C:30 T:0 | | | | | | | | | | | | | | | |
| | JPHHDCAP_3 | 42008 | snp | A | T | T:20 A:0 | CDS | + | 72/303 | 24/100 | synonymous_variant c.72A>T p.Ala24Ala | JPHHDCAP_03198 | HigA1 | | | | | | | | |
| | JPHHDCAP_4 | 76996 | snp | C | T | T:20 C:0 | CDS | - | 238/1281 | 80/426 | missense_variant c.238G>A p.Asp80Asn | JPHHDCAP_03293 | CzcD | | | | | | | | |
| | JPHHDCAP_4 | 77095 | snp | T | C | C:20 T:0 | CDS | - | 139/1281 | 47/426 | missense_variant c.139A>G p.Lys47Glu | JPHHDCAP_03293 | CzcD | | | | | | | | |
| | JPHHDCAP_4 | 77302 | complex | ATC | TTT | TTT:20 ATC:0 | | | | | | | | | | | | | | | |
| | JPHHDCAP_4 | 77316 | snp | T | C | C:20 T:0 | | | | | | | | | | | | | | | |
| | JPHHDCAP_4 | 77415 | snp | G | C | C:20 G:0 | CDS | + | 93/393 | 31/130 | synonymous_variant c.93G>C p.Pro31Pro | JPHHDCAP_03294 | MerR | | | | | | | | |
| | JPHHDCAP_4 | 77449 | snp | G | A | A:20 G:0 | CDS | + | 127/393 | 43/130 | missense_variant c.127G>A p.Val43Ile | JPHHDCAP_03294 | MerR | | | | | | | | |
| | JPHHDCAP_4 | 77472 | snp | A | G | G:20 A:0 | CDS | + | 150/393 | 50/130 | synonymous_variant c.150A>G p.Lys50Lys | JPHHDCAP_03294 | MerR | | | | | | | | |
| Gen_AL28 WT1 | JPHHDCAP_4 | 77698 | snp | T | C | C:15 T:0 | CDS | + | 376/393 | 126/130 | synonymous_variant c.376T>C p.Leu126Leu | JPHHDCAP_03294 | MerR | 0.25 | 0.06 | 0.25 | 8 | 0.06 | 0.03 | <0.015 | <0.12 |
| | JPHHDCAP_1 | 41290 | snp | A | C | C:20 A:0 | | | | | | | | | | | | | | | |
| | JPHHDCAP_2 | 405216 | snp | G | A | A:20 G:0 | CDS | + | 337/1005 | 113/334 | missense_variant c.337G>A p.Gly113Ser | JPHHDCAP_00497 | Ribokinase | | | | | | | | |
| | JPHHDCAP_3 | 8043 | snp | A | C | C:20 A:0 | CDS | - | 409/2463 | 137/820 | missense_variant c.409T>G p.Leu137Val | JPHHDCAP_03157 | VirB4 | | | | | | | | |
| | JPHHDCAP_3 | 21503 | snp | C | A | A:20 C:0 | CDS | + | 630/777 | 210/258 | missense_variant c.630C>A p.Ser210Arg | JPHHDCAP_03170 | Soj2 | | | | | | | | |
| | JPHHDCAP_3 | 21513 | snp | G | T | T:20 G:0 | CDS | + | 640/777 | 214/258 | missense_variant c.640G>T p.Val214Phe | JPHHDCAP_03170 | Soj2 | | | | | | | | |
| | JPHHDCAP_3 | 22311 | snp | G | T | T:20 G:0 | CDS | + | 658/1038 | 220/345 | stop_gained c.658G>T p.Glu220* | JPHHDCAP_03171 | KorB | | | | | | | | |
| | JPHHDCAP_3 | 22783 | snp | G | T | T:18 G:0 | | | | | | | | | | | | | | | |
| | JPHHDCAP_3 | 28803 | snp | C | G | G:20 C:0 | CDS | + | 189/426 | 63/141 | synonymous_variant c.189C>G p.Leu63Leu | JPHHDCAP_03181 | MerC | | | | | | | | |
| | JPHHDCAP_3 | 37827 | snp | A | G | G:10 A:0 | CDS | + | 514/600 | 172/199 | missense_variant c.514A>G p.Arg172Gly | JPHHDCAP_03193 | Hin2 | | | | | | | | |
| | JPHHDCAP_3 | 41687 | snp | T | C | C:31 T:0 | | | | | | | | | | | | | | | |
| | JPHHDCAP_3 | 42008 | snp | A | T | T:20 A:0 | CDS | + | 72/303 | 24/100 | synonymous_variant c.72A>T p.Ala24Ala | JPHHDCAP_03198 | HigA1 | | | | | | | | |
| | JPHHDCAP_3 | 45049 | snp | G | T | T:20 G:0 | | | | | | | | | | | | | | | |
| | JPHHDCAP_3 | 51801 | snp | T | G | G:20 T:0 | CDS | - | 488/1278 | 163/425 | missense_variant c.488A>C p.Tyr163Ser | JPHHDCAP_03207 | hypothetical protein | | | | | | | | |
| Gen_AL28 WT2 | JPHHDCAP_4 | 76996 | snp | C | T | T:12 C:0 | CDS | - | 238/1281 | 80/426 | missense_variant c.238G>A p.Asp80Asn | JPHHDCAP_03293 | CzcD | 0.12 | <0.03 | 0.25 | 8 | 0.06 | 0.03 | <0.015 | <0.12 |
| | JPHHDCAP_1 | 62161 | snp | T | C | C:20 T:0 | | | | | | | | | | | | | | | |
| | JPHHDCAP_3 | 10760 | snp | T | C | C:20 T:0 | CDS | - | 1395/2253 | 465/750 | synonymous_variant c.1395A>G p.Glu465Glu | JPHHDCAP_03160 | TopA2 | | | | | | | | |
| | JPHHDCAP_3 | 10799 | snp | T | A | A:20 T:0 | CDS | - | 1356/2253 | 452/750 | synonymous_variant c.1356A>T p.Pro452Pro | JPHHDCAP_03160 | TopA2 | | | | | | | | |
| | JPHHDCAP_3 | 10827 | snp | T | A | A:20 T:0 | CDS | - | 1328/2253 | 443/750 | missense_variant c.1328A>T p.Glu443Val | JPHHDCAP_03160 | TopA2 | | | | | | | | |
| | JPHHDCAP_3 | 11645 | snp | T | A | A:16 T:0 | CDS | - | 510/2253 | 170/750 | synonymous_variant c.510A>T p.Ser170Ser | JPHHDCAP_03160 | TopA2 | | | | | | | | |
| | JPHHDCAP_3 | 15916 | snp | A | T | T:20 A:0 | CDS | + | 674/1191 | 225/396 | missense_variant c.674A>T p.Glu225Val | JPHHDCAP_03165 | hypothetical protein | | | | | | | | |
| | JPHHDCAP_3 | 22781 | snp | A | C | C:13 A:0 | | | | | | | | | | | | | | | |
| | JPHHDCAP_3 | 35260 | snp | G | T | T:20 G:0 | CDS | - | 492/1041 | 164/346 | missense_variant c.492C>A p.Phe164Leu | JPHHDCAP_03189 | Acr3 | | | | | | | | |
| | JPHHDCAP_3 | 37827 | snp | A | G | G:10 A:0 | CDS | + | 514/600 | 172/199 | missense_variant c.514A>G p.Arg172Gly | JPHHDCAP_03193 | Hin2 | | | | | | | | |
| | JPHHDCAP_3 | 40361 | snp | T | G | G:20 T:0 | CDS | + | 759/1350 | 253/449 | missense_variant c.759T>G p.Ser253Arg | JPHHDCAP_03196 | ChrA1 | | | | | | | | |
| | JPHHDCAP_3 | 41687 | snp | T | C | C:32 T:0 | | | | | | | | | | | | | | | |
| | JPHHDCAP_3 | 41695 | snp | A | G | G:32 A:0 | | | | | | | | | | | | | | | |
| | JPHHDCAP_3 | 41710 | snp | T | C | C:34 T:0 | | | | | | | | | | | | | | | |
| | JPHHDCAP_3 | 41764 | snp | T | C | C:33 T:0 | CDS | + | 15/195 | May-64 | synonymous_variant c.15T>C p.Arg5Arg | JPHHDCAP_03197 | hypothetical protein | | | | | | | | |
| | JPHHDCAP_3 | 42008 | snp | A | T | T:20 A:0 | CDS | + | 72/303 | 24/100 | synonymous_variant c.72A>T p.Ala24Ala | JPHHDCAP_03198 | HigA1 | | | | | | | | |
| | JPHHDCAP_3 | 43869 | snp | C | A | A:20 C:0 | CDS | + | 1371/1878 | 457/625 | missense_variant c.1371C>A p.Asp457Glu | JPHHDCAP_03199 | Kup2 | | | | | | | | |
| | JPHHDCAP_3 | 45003 | snp | T | G | G:20 T:0 | | | | | | | | | | | | | | | |
| | JPHHDCAP_3 | 47250 | snp | T | G | G:11 T:0 | CDS | + | 619/1446 | 207/481 | missense_variant c.619T>G p.Phe207Val | JPHHDCAP_03202 | DNA-cytosine methyltransferase | | | | | | | | |
| | JPHHDCAP_3 | 54268 | snp | T | A | A:20 T:0 | CDS | - | 570/2247 | 190/748 | synonymous_variant c.570A>T p.Pro190Pro | JPHHDCAP_03209 | hypothetical protein | | | | | | | | |
| Gen_AL28 M1 | JPHHDCAP_3 | 54400 | snp | A | C | C:20 A:0 | CDS | - | 438/2247 | 146/748 | synonymous_variant c.438T>G p.Gly146Gly | JPHHDCAP_03209 | hypothetical protein | 0.25 | <0.03 | 0.25 | 8 | 0.25 | 0.03 | 0.03 | <0.12 |
| | JPHHDCAP_1 | 62161 | snp | T | C | C:20 T:0 | | | | | | | | | | | | | | | |
| | JPHHDCAP_2 | 405216 | snp | G | A | A:20 G:0 | CDS | + | 337/1005 | 113/334 | missense_variant c.337G>A p.Gly113Ser | JPHHDCAP_00497 | Ribokinase | | | | | | | | |
| | JPHHDCAP_3 | 1505 | del | AT | A | A:19 AT:0 | CDS | - | 540/1425 | 180/474 | frameshift_variant c.540delA p.Gln180fs | JPHHDCAP_03151 | hypothetical protein | | | | | | | | |
| | JPHHDCAP_3 | 13375 | snp | A | T | T:12 A:0 | CDS | + | 195/246 | 65/81 | missense_variant c.195A>T p.Lys65Asn | JPHHDCAP_03162 | hypothetical protein | | | | | | | | |
| | JPHHDCAP_3 | 13392 | del | TTTATAAACAATTACAAGAA | T | T:10 TTTATA | CDS | + | 213/246 | 71/81 | frameshift_variant&stop_lost&splice_region_variant c.213_*15delTTATAAA( | JPHHDCAP_03162 | hypothetical protein | | | | | | | | |
| | JPHHDCAP_3 | 13487 | snp | C | G | G:14 C:0 | CDS | - | 286/321 | 96/106 | missense_variant c.286G>C p.Asp96His | JPHHDCAP_03163 | hypothetical protein | | | | | | | | |
| | JPHHDCAP_3 | 18247 | snp | G | T | T:20 G:0 | | | | | | | | | | | | | | | |
| | JPHHDCAP_3 | 20068 | snp | C | A | A:20 C:0 | | | | | | | | | | | | | | | |
| | JPHHDCAP_3 | 25339 | snp | A | G | G:20 A:0 | CDS | - | 305/936 | 102/311 | missense_variant c.305T>C p.Ile102Thr | JPHHDCAP_03174 | hypothetical protein | | | | | | | | |
| | JPHHDCAP_3 | 28771 | snp | C | A | A:18 C:0 | CDS | + | 157/426 | 53/141 | missense_variant c.157C>A p.Leu53Met | JPHHDCAP_03181 | MerC | | | | | | | | |
| | JPHHDCAP_3 | 28820 | snp | G | T | T:18 G:0 | CDS | + | 206/426 | 69/141 | missense_variant c.206G>T p.Trp69Leu | JPHHDCAP_03181 | MerC | | | | | | | | |
| | JPHHDCAP_3 | 28992 | del | TCGCCGCTGCGGGCCGGAC | T | T:16 TCGCCG | CDS | + | 380/426 | 127/141 | frameshift_variant c.380_399delGCCGCTGCGGGCCGGACAGC p.Arg127fs | JPHHDCAP_03181 | MerC | | | | | | | | |
| | JPHHDCAP_3 | 29092 | snp | A | G | G:19 A:0 | CDS | + | 14/1686 | 5/561 | missense_variant c.14A>G p.Lys5Arg | JPHHDCAP_03182 | MerA | | | | | | | | |
| Gen_AL28 M2 | JPHHDCAP_3 | 41687 | snp | T | C | C:21 T:0 | | | | | | | | 0.25 | <0.03 | 0.25 | 8 | 0.25 | 0.03 | 0.03 | <0.12 |
| | JPHHDCAP_3 | 41687 | snp | T | C | C:26 T:0 | | | | | | | | | | | | | | | |
| | JPHHDCAP_3 | 42008 | snp | A | T | T:20 A:0 | CDS | + | 72/303 | 24/100 | synonymous_variant c.72A>T p.Ala24Ala | JPHHDCAP_03198 | HigA1 | | | | | | | | |
| Gen_AL28 M3 | JPHHDCAP_4 | 76996 | snp | C | T | T:12 C:0 | CDS | - | 238/1281 | 80/426 | missense_variant c.238G>A p.Asp80Asn | JPHHDCAP_03293 | CzcD | 0.25 | <0.03 | 0.25 | 8 | 0.25 | 0.03 | 0.03 | <0.12 |
| | JPHHDCAP_3 | 6887 | snp | T | G | G:20 T:0 | CDS | - | 1565/2463 | 522/820 | missense_variant c.1565A>C p.Asn522Thr | JPHHDCAP_03157 | VirB4 | | | | | | | | |
| | JPHHDCAP_3 | 23369 | snp | G | C | C:20 G:0 | CDS | + | 439/669 | 147/222 | missense_variant c.439G>C p.Val147Leu | JPHHDCAP_03172 | hypothetical protein | | | | | | | | |
| | JPHHDCAP_3 | 37827 | snp | A | G | G:10 A:0 | CDS | + | 514/600 | 172/199 | missense_variant c.514A>G p.Arg172Gly | JPHHDCAP_03193 | Hin2 | | | | | | | | |
| | JPHHDCAP_3 | 41687 | snp | T | C | C:33 T:0 | | | | | | | | | | | | | | | |
| | JPHHDCAP_3 | 42008 | snp | A | T | T:20 A:0 | CDS | + | 72/303 | 24/100 | synonymous_variant c.72A>T p.Ala24Ala | JPHHDCAP_03198 | HigA1 | | | | | | | | |
| | JPHHDCAP_4 | 76996 | snp | C | T | T:20 C:0 | CDS | - | 238/1281 | 80/426 | missense_variant c.238G>A p.Asp80Asn | JPHHDCAP_03293 | CzcD | | | | | | | | |
| | JPHHDCAP_4 | 77095 | snp | T | C | C:20 T:0 | CDS | - | 139/1281 | 47/426 | missense_variant c.139A>G p.Lys47Glu | JPHHDCAP_03293 | CzcD | | | | | | | | |
| | JPHHDCAP_4 | 77121 | snp | A | G | G:20 A:0 | CDS | - | 113/1281 | 38/426 | missense_variant c.113T>C p.Leu38Pro | JPHHDCAP_03293 | CzcD | | | | | | | | |
| | JPHHDCAP_4 | 77148 | snp | T | C | C:20 T:0 | CDS | - | 86/1281 | 29/426 | missense_variant c.86A>G p.Asn29Ser | JPHHDCAP_03293 | CzcD | | | | | | | | |
| | JPHHDCAP_4 | 77225 | snp | A | G | G:20 A:0 | CDS | - | 9/1281 | 3/426 | synonymous_variant c.9T>C p.Gly3Gly | JPHHDCAP_03293 | CzcD | | | | | | | | |
| | JPHHDCAP_4 | 77235 | snp | A | G | G:20 A:0 | | | | | | | | | | | | | | | |
| | JPHHDCAP_4 | 77253 | ins | A | AT | AT:19 A:0 | | | | | | | | | | | | | | | |
| | JPHHDCAP_4 | 77302 | complex | ATC | TTT | TTT:20 ATC:0 | | | | | | | | | | | | | | | |
| | JPHHDCAP_4 | 77316 | snp | T | C | C:20 T:0 | | | | | | | | | | | | | | | |
| | JPHHDCAP_4 | 77544 | complex | ACCTG | TCCTA | TCCTA:16 AC | CDS | + | 222/393 | 74/130 | missense_variant c.222_226delACCTGinsTCCTA p.LysProGlu74AsnProLys | JPHHDCAP_03294 | MerR | | | | | | | | |
| | JPHHDCAP_4 | 77641 | snp | C | T | T:15 C:0 | CDS | + | 319/393 | 107/130 | synonymous_variant c.319C>T p.Leu107Leu | JPHHDCAP_03294 | MerR | | | | | | | | |
| Gen_AL28 M4 | JPHHDCAP_4 | 77698 | snp | T | C | C:15 T:0 | CDS | + | 376/393 | 126/130 | synonymous_variant c.376T>C p.Leu126Leu | JPHHDCAP_03294 | MerR | 0.25 | <0.03 | 0.25 | 8 | 0.25 | 0.03 | 0.03 | <0.12 |
| | JPHHDCAP_1 | 41290 | snp | A | C | C:20 A:0 | | | | | | | | | | | | | | | |

| | | | | | | | | | | | | | | | | | | | | | |
|---|---|---|---|---|---|---|---|---|---|---|---|---|---|---|---|---|---|---|---|---|---|
| | JPHHDCAP_2 | 917216 | snp | G | A | A:20 G:0 | CDS | - | 562/1899 | 188/632 | missense_variant c.562C>T p.Leu188Phe | JPHHDCAP_01023 | SdhA | | | | | | | | |
| | JPHHDCAP_2 | 1075139 | snp | C | T | T:20 C:0 | CDS | - | 961/1014 | 321/337 | missense_variant c.961G>A p.Ala321Thr | JPHHDCAP_01157 | hypothetical protein | | | | | | | | |
| | JPHHDCAP_3 | 19336 | snp | T | G | G:20 T:0 | CDS | + | 279/519 | 93/172 | missense_variant c.279T>G p.Asp93Glu | JPHHDCAP_03168 | hypothetical protein | | | | | | | | |
| | JPHHDCAP_3 | 21513 | snp | G | T | T:20 G:0 | CDS | + | 640/777 | 214/258 | missense_variant c.640G>T p.Val214Phe | JPHHDCAP_03170 | Soj2 | | | | | | | | |
| | JPHHDCAP_3 | 23082 | complex | CAGATTATTCAGG | AATATAATT/ | AATATAATT/ | CDS | + | 152/669 | 51/222 | stop_gained c.152_164delCAGATTATTCAGGinsAATATAATTAAGT p.ProAspTy | JPHHDCAP_03172 | hypothetical protein | | | | | | | | |
| | JPHHDCAP_3 | 24888 | snp | C | A | A:20 C:0 | CDS | - | 756/936 | 252/311 | synonymous_variant c.756G>T p.Ser252Ser | JPHHDCAP_03174 | hypothetical protein | | | | | | | | |
| | JPHHDCAP_3 | 28127 | snp | T | G | G:20 T:0 | CDS | + | 180/351 | 60/116 | synonymous_variant c.180T>G p.Val60Val | JPHHDCAP_03179 | MerT | | | | | | | | |
| | JPHHDCAP_3 | 30540 | del | AAGCTGGTCATCG | A | A:18 AAGCT( | CDS | + | 1465/1686 | 489/561 | conservative_inframe_deletion c.1465_1476delCTGGTCATCGAG p.Leu489_G | JPHHDCAP_03182 | MerA | | | | | | | | |
| | JPHHDCAP_3 | 31044 | del | TGCGGCAACTCGTCGAGCG | T | T:16 TGCGG( | CDS | + | 267/366 | 89/121 | frameshift_variant c.267_289delGCAACTCGTCGAGCGTCGGCGCG p.Gln90fs | JPHHDCAP_03183 | hypothetical protein | | | | | | | | |
| | JPHHDCAP_3 | 34129 | snp | A | T | T:20 A:0 | CDS | - | 578/705 | 193/234 | missense_variant c.578T>A p.Val193Glu | JPHHDCAP_03188 | ArsH | | | | | | | | |
| | JPHHDCAP_3 | 34238 | snp | T | A | A:20 T:0 | CDS | - | 469/705 | 157/234 | missense_variant c.469A>T p.Met157Leu | JPHHDCAP_03188 | ArsH | | | | | | | | |
| | JPHHDCAP_3 | 37827 | snp | A | G | G:10 A:0 | CDS | + | 514/600 | 172/199 | missense_variant c.514A>G p.Arg172Gly | JPHHDCAP_03193 | Hin2 | | | | | | | | |
| | JPHHDCAP_3 | 41687 | snp | T | C | C:32 T:0 | | | | | | | | | | | | | | | |
| Gen_AL28 M5 | JPHHDCAP_3 | 42008 | snp | A | T | T:20 A:0 | CDS | + | 72/303 | 24/100 | synonymous_variant c.72A>T p.Ala24Ala | JPHHDCAP_03198 | HigA1 | 0.12 | <0.03 | 0.25 | 8 | 0.06 | 0.015 | <0.015 | 0.25 |
