## Supplementary material for "Differential development of antibiotic resistance and virulence between *Acinetobacter* species": S1-4_S6_S12_S14-19

**Supplementary File 1**

**Table** **S1** Strains used in this study

| **Strain identifier** | **Associated Name** | **Reference** |
| --- | --- | --- |
| AB05 | *A. baumannii* AYE | (1) |
| AB06 | *A. baumannii* UKA2 | (2) |
| AB07 | *A. baumannii* NCTC 13302 | UKHSA |
| AB17 | *A. baumannii* MPT156 | This study |
| AB18 | *A. baumannii* All8 | This study |
| AB19 | *A. baumannii* W1 | This study |
| AB20 | *A. baumannii* NCTC 10303 | (3) |
| AB22 | *A. baumannii* NCTC 13423 | (4) |
| AB23 | *A. baumannii* NCTC 13424 | (5) |
| AB25 | *A. baumannii* UKA2b | This study |
| AB27 | *A. baumannii* UKA15 | This study |
| AL04 | *A. lwoffii* clinical isolate | This study |
| AL28 | *A. lwoffii* NCTC 5867 | UKHSA |
| AL29 | *A. lwoffii* NCTC 10309 | (3) |
| AL31 | *A. lwoffii* NCTC 5866 | (6) |
| AL32 | *A. lwoffii* MW156 | This study |
| AL33 | *A. lwoffii* MW158 | This study |
| AL44 | *A. lwoffii* E9 | This study |
| AL51 | *A. lwoffii* agricultural isolate | This study |
| AL52 | *A. lwoffii* agricultural isolate | This study |
| AL54 | *A. lwoffii* agricultural isolate | This study |
| AL55 | *A. lwoffii* agricultural isolate | This study |

**Text** **S2**

Biofilm. Formation in static conditions and under laminar flow

Biofilm mass formed in static conditions was measured using a crystal violet biofilm assay as previously described in Ref. (7).

Biofilm formation under laminar flow was measured using the Bioflux system as previously described (8). After 48 hours, flow cells were pulsed at 5 dyne/cm^2^ for 5s to capture images of attached biofilm only. Experiments were performed with two technical and two biological replicates. Images were analysed using ImageJ (9), where the percentage of black pixels in the field of view was calculated as a proxy for biofilm growth. The median percentage coverage was then plotted on Graphpad Prism (10).

Antimicrobial susceptibility of established *A. baumannii* and *A. lwoffii* biofilms

The minimum biofilm eradication concentration of established static biofilms was determined using a peg lid model as previously described (11). Compounds tested included cefotaxime (Fisher #10084487), chlorhexidine (Sigma #C9394), ciprofloxacin (Fisher #13531640), meropenem (TCI Chemicals #M2279), oxacillin (Sigma, #O1002), tetracycline (Fisher #10460264), triclosan (Sigma, #72779) and rifampicin (Fisher #10533325).

**Table** **S3** Whole genome sequences generated in this study. All raw reads are listed on NCBI under PRJNA918592.

| **Isolate identifier** | **BioSample Accession** |
| --- | --- |
| AL04 | SAMN32597906 |
| AB06 | SAMN32597907 |
| AB07 | SAMN32597908 |
| AB17 | SAMN32597909 |
| AB18 | SAMN32597910 |
| AB19 | SAMN32597911 |
| AB20 | SAMN32597912 |
| AB22 | SAMN32597913 |
| AB23 | SAMN32597914 |
| AB25 | SAMN32597915 |
| AB27 | SAMN32597916 |
| AL29 | SAMN32597917 |
| AL31 | SAMN32597918 |
| AL32 | SAMN32597919 |
| AL33 | SAMN32597920 |
| AL44 | SAMN32597921 |
| AL51 | SAMN32597922 |
| AL52 | SAMN32597923 |
| AL54 | SAMN32597924 |
| AL55 | SAMN32597925 |
| Meropenem Evolution: AB18 WT1 | SAMN32597926 |
| Meropenem Evolution: AB18 WT2 | SAMN32597927 |
| Meropenem Evolution: AB18 Mutant 1 | SAMN32597928 |
| Meropenem Evolution: AB18 Mutant 2 | SAMN32597929 |
| Meropenem Evolution: AB18 Mutant 3 | SAMN32597930 |
| Meropenem Evolution: AB18 Mutant 4 | SAMN32597931 |
| Meropenem Evolution: AB18 Mutant 5 | SAMN32597932 |
| Meropenem Evolution AL28 WT 1 | SAMN32597933 |
| Meropenem Evolution AL28 WT 2 | SAMN32597934 |
| Meropenem Evolution: AL28 Mutant 1 | SAMN32597935 |
| Meropenem Evolution: AL28 Mutant 2 | SAMN32597936 |
| Meropenem Evolution: AL28 Mutant 3 | SAMN32597937 |
| Meropenem Evolution: AL28 Mutant 4 | SAMN32597938 |
| Meropenem Evolution: AL28 Mutant 5 | SAMN32597939 |
| Ciprofloxacin Evolution: AB18 WT1 | SAMN35816247 |
| Ciprofloxacin Evolution: AB18 WT2 | SAMN35816248 |
| Ciprofloxacin Evolution: AB18 Mutant 1 | SAMN35816249 |
| Ciprofloxacin Evolution: AB18 Mutant 2 | SAMN35816250 |
| Ciprofloxacin Evolution: AB18 Mutant 3 | SAMN35816251 |
| Ciprofloxacin Evolution: AB18 Mutant 4 | SAMN35816252 |
| Ciprofloxacin Evolution: AB18 Mutant 5 | SAMN35816253 |
| Ciprofloxacin Evolution AL28 WT 1 | SAMN35816254 |
| Ciprofloxacin Evolution AL28 WT 2 | SAMN35816255 |
| Ciprofloxacin Evolution: AL28 Mutant 1 | SAMN35816256 |
| Ciprofloxacin Evolution: AL28 Mutant 2 | SAMN35816257 |
| Ciprofloxacin Evolution: AL28 Mutant 3 | SAMN35816258 |
| Ciprofloxacin Evolution: AL28 Mutant 4 | SAMN35816259 |
| Ciprofloxacin Evolution: AL28 Mutant 5 | SAMN35816260 |
| Gentamicin Evolution: AB18 WT1 | SAMN35816261 |
| Gentamicin Evolution: AB18 WT2 | SAMN35816262 |
| Gentamicin Evolution: AB18 Mutant 1 | SAMN35816263 |
| Gentamicin Evolution: AB18 Mutant 2 | SAMN35816264 |
| Gentamicin Evolution: AB18 Mutant 3 | SAMN35816265 |
| Gentamicin Evolution: AB18 Mutant 4 | SAMN35816266 |
| Gentamicin Evolution: AB18 Mutant 5 | SAMN35816267 |
| Gentamicin Evolution AL28 WT 1 | SAMN35816268 |
| Gentamicin Evolution AL28 WT 2 | SAMN35816269 |
| Gentamicin Evolution: AL28 Mutant 1 | SAMN35816270 |
| Gentamicin Evolution: AL28 Mutant 2 | SAMN35816271 |
| Gentamicin Evolution: AL28 Mutant 3 | SAMN35816272 |
| Gentamicin Evolution: AL28 Mutant 4 | SAMN35816273 |
| Gentamicin Evolution: AL28 Mutant 5 | SAMN35816274 |

AL = *A. lwoffii* AB= *A. baumannii*

**Text S4**

Measurement of twitching motility

The twitching capacity of *A. baumannii* and *A. lwoffii* was determined as detailed previously (12). Sub-surface twitching halos were stained and measured using crystal violet.

Comparing the growth of *A. baumannii* and *A. lwoffii*

Growth in LB and male AB human serum (Merck, #H4522) was measured for 16 and 24 hours in a Fluostar Omega plate reader (BMG LabTech). The OD_600_ was measured at ten-minute intervals and a growth curve was plotted on Prism (v.9, Graphpad) (10). Mean generation time was computed using R package growthcurver (13) and comparative statistics performed in Prism. Human serum was used both fresh (NHS - normal human serum) and heat inactivated at 56°C for 1 hour (HIS - heat inactivated serum).

Survival in serum was also quantified. Briefly, cells were grown to mid-log phase and then diluted to 1x10^6^ CFU/mL. 10 μL of cells were added to 90 μL of either NHS or LB. Cells were incubated at 37°C with gentle rocking (20 rpm). 10 μL was removed at 0 minutes, 45 minutes, 1.5 hours, 3 hour and 24 hours and serially diluted to enumerate colony forming units (CFUs).

Strains were also grown in 5 mL synthetic wound fluid at 37°C with shaking for five hours and diluted to an OD of ~0.1. Wound fluid was created using a previously published method (14) in 24 well plates and the bacteria were statically incubated with humidity for 3 days at 37°C and visible growth was compared.

**Fig. S6** A phylogenetic tree of MAFFT aligned *rep* genes from (15) and additional *A. lwoffii rep* genes from available complete genome sequences on NCBI (red). Arrows point to zoomed in areas of the tree to show additional *A. lwoffii rep* genes

**Fig. S12** *A. lwoffii* forms less biofilm in static conditions than *A. baumannii,* but both species form similar levels of biofilm under laminar flow. a- total biofilm density grouped by species, b- individual strain biofilm formation. Median values of three biological repeats plotted, along with interquartile range. Two-tailed Welch’s T-test proved significance when comparing the species, p <0.0001. c- percentage coverage of biofilm formed on a flow cell under laminar flow in the Bioflux model. Median coverage is plotted with interquartile range. d- number of biofilm associated genes found in whole genome sequences, whiskers show minimum and maximum values. Welch’s T test was used to compare the number of genes in either species after a random permutation test accounted for different sample sizes, p <0.0001. Pink - *A. baumannii*, yellow - *A. lwoffii*.

**Fig. S14** Scanning electron micrographs of *A. baumannii* (AB) and *A. lwoffii* (AL) strains taken on an Apreo 2 (Thermo Fisher).

**Table S15** Mean generation times (hours) for strains in LB at different temperatures calculated using Growthcurver.

| **Strain** | **37°C – Mean generation time ± standard error** | **30°C - Mean generation time ± standard error** | **25°C - Mean generation time ± standard error** |
| --- | --- | --- | --- |
| **AL04** | 1.070 ± 0.165 | 0.515 ± 0.058 | 0.960 ± 0.170 |
| **AB05** | 0.650 ± 0.003 | 0.758 ± 0.070 | 1.002 ± 0.016 |
| **AB19** | 0.595 ± 0.005 | 0.758 ± 0.067 | 0.813 ± 0.032 |
| **AB20** | 0.563 ± 0.017 | 0.703 ± 0.044 | 0.839 ± 0.019 |
| **AB25** | 0.861 ± 0.014 | 0.880 ± 0.063 | 0.972 ± 0.012 |
| **AB27** | 0.701 ± 0.006 | 0.740 ± 0.058 | 0.918 ± 0.045 |
| **AL28** | 0.797 ± 0.406 | 0.406 ± 0.032 | 0.818 ± 0.195 |
| **AL29** | 1.689 ± 0.448 | 0.832 ± 0.189 | 1.362 ± 0.086 |
| **AL32** | 0.348 ± 0.033 | 0.337 ± 0.076 | 0.637 ± 0.135 |
| **AL33** | 0.434 ± 0.117 | 0.648 ± 0.183 | 0.784 ± 0.156 |

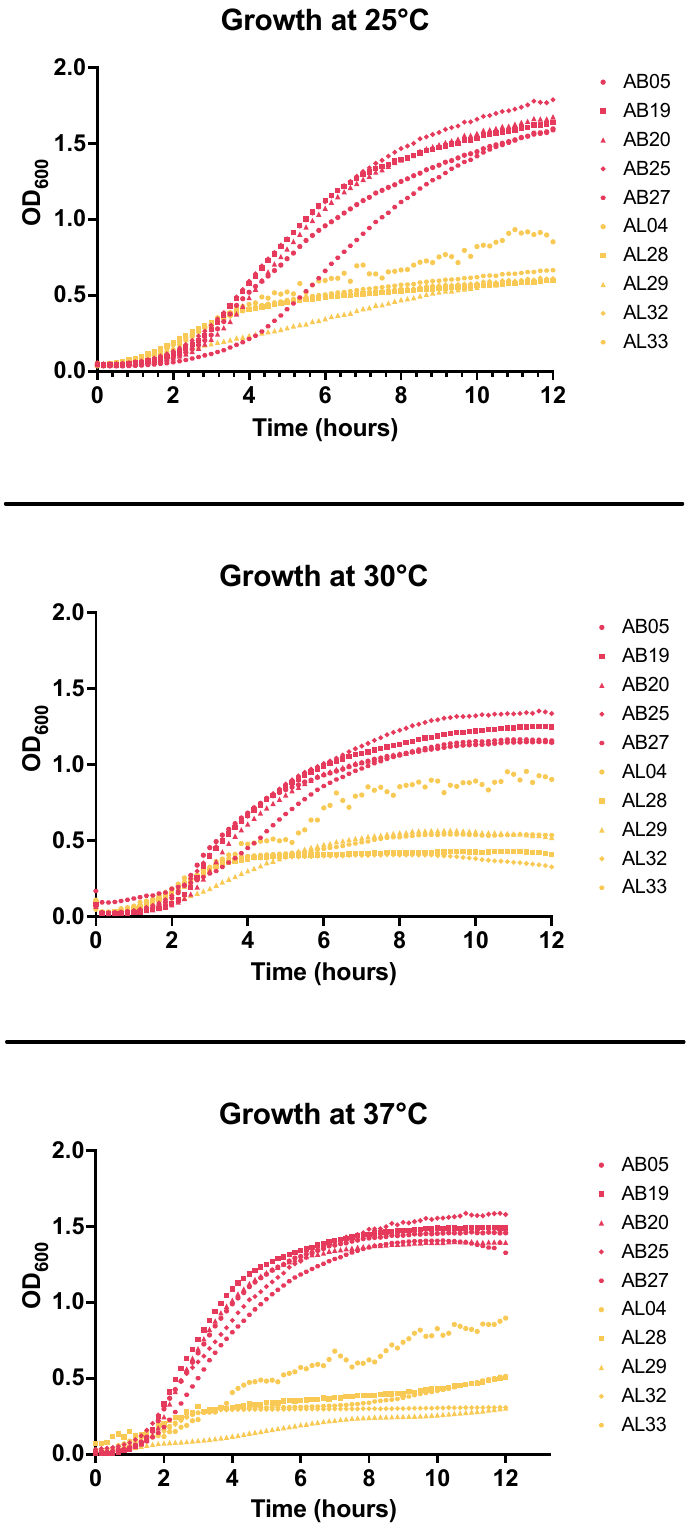

**Fig S16** Growth curves of *A. baumannii* (pink)and *A. lwoffii* (yellow) in LB over twelve hours at 37°C. Optical density at 600_nm_ was measured every twenty minutes in a FLUOstar Omega Plate Reader.

**Table S17:** Mean generation times (hours) of strains grown in LB and human serum at 37°C.

| **Strain** | **LB** | **Heat-Inactivated Serum** | **Normal Human Serum** |
| --- | --- | --- | --- |
|  | **Mean generation time ± standard error** | **Mean generation time ± standard error** | **Mean generation time ± standard error** |
| **AL04** | 0.926 ± 0.26 | 3.257 ± 1.284 | 4.611 ± 1.728 |
| **AB05** | 0.910 ± 0.088 | 2.507 ± 0.04 | 3.972 ± 1.62 |
| **AB18** | 0.691 ± 0.067 | 1.385 ± 0.012 | 1.495 ± 0.12 |
| **AL28** | 0.492 ± 0.066 | * | * |

* = due to the lack of growth of AL28 in serum it was not possible to compute accurate mean generation times, see supplementary Figure S15.

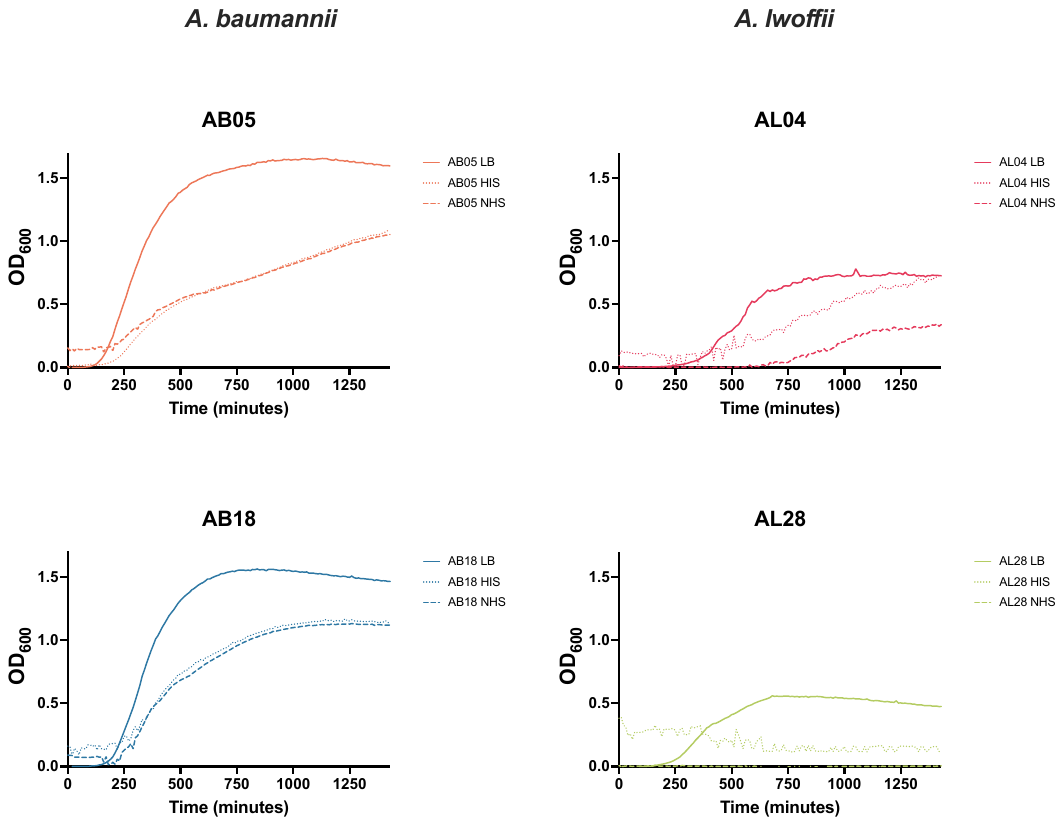

**Fig S18** The growth of *A. baumannii* and *A. lwoffii* in human serum

NHS = normal human serum, HIS = heat inactivated serum

**Table S19** Growth of *A. baumannii* (AB05, AB18) and *A. lwoffii* (AL04, AL28) in a synthetic wound model after three days incubation at 37°C

| **Strain** | **Day 1** | **Day 2** | **Day 3** |
| --- | --- | --- | --- |
| AL04 | - | - | - |
| AL28 | - | - | - |
| AB05 | - | + | ++ |
| AB18 | + | ++ | ++ |
| Uninoculated wound | - | - | - |
| Sterile synthetic wound fluid | - | - | - |
